# Complex IV Subunit C15orf48 Integrates Electron Transport Chain Function with Amino Acid and Lipid Metabolism to Regulate Innate Immunity

**DOI:** 10.64898/2026.11.26.769465

**Authors:** Cheryl QE Lee, Sze Huey Leong, Sheryl Beh, Kimberle Shen, Chengwei Zhong, Radiance Lim, Yunxia Liu, Federico Torta, Guo-Fang Zhang, Yibin Wang, Jacques Behmoaras, Christopher B. Newgard, Jin Miao Chen, Lena Ho

**Affiliations:** Signature Research Program in Cardiovascular and Metabolic Disorders, Duke-National University of Singapore Medical School, Singapore 169857; Infectious Diseases Translational Research Programme, Department of Microbiology and Immunology, Yong Loo Lin School of Medicine, National University of Singapore, 5 Science Drive 2, Singapore 117545; Genome Institute of Singapore, A*STAR, 60 Biopolis Street, Singapore 138672; Bioinformatics Institute, A*STAR, 30 Biopolis Street, Singapore 138672; Singapore Lipidomics Incubator, Life Sciences Institute, Singapore 117456; Precision Medicine Translational Research Programme and Department of Biochemistry, Yong Loo Lin School of Medicine, National University of Singapore, Singapore 117597; Sarah W. Stedman Nutrition and Metabolism Center & Duke Molecular Physiology Institute, Duke University, Durham, NC 27701, USA; Department of Medicine, Division of Endocrinology, Metabolism and Nutrition, Duke University Medical Center, Durham, NC 27701, USA; National Heart Centre Singapore, Singapore 169609; Centre for Computational Biology, Duke-National University of Singapore Medical School, Singapore 169857; Centre for Biomedical Data Science and Program in Cancer and Stem Cell Biology, Duke-NUS Medical School, Singapore; Immunology Translational Research Program, Department of Microbiology and Immunology, Yong Loo Lin School of Medicine, National University of Singapore, 5 Science Drive 2, Singapore 117545

## Abstract

Despite extensive recognition of immunometabolism in innate immune regulation, how mitochondrial bioenergetic and metabolic pathways in monocytes interface to control inflammatory responses remains poorly understood. Here, we uncover that *C15orf48* (MOCCI/NMES1/COXFA4L3), an inflammation-associated mitochondrial complex IV (CIV) subunit, orchestrates electron transport chain (ETC) function, fatty acid oxidation (FAO), and branched-chain amino acid (BCAA) catabolism to control monocyte activation. Loss of *C15orf48* globally and in monocytes confers host tolerance and resistance to bacterial pneumonia and LPS-induced endotoxemia, characterized by reduced inflammatory gene expression, cytokine secretion, and monocyte activation. Mechanistically, *C15orf48* increases mitochondrial ROS production and maintains NAD pools, which in turn supports long-chain FAO and BCAA catabolism required for robust monocyte activation and inflammatory cytokine production. These findings define a mitochondrial CIV metabolic axis directed by *C15orf48* as central to monocyte activation and innate immunity, filling a key knowledge gap in immunometabolic control and presenting new targets for therapy in infection and sepsis.

## Introduction

Uncontrolled systemic inflammation is the primary driver of morbidity and mortality in sepsis, characterized by a dysregulated immune response and hypercytokinemia, commonly referred to as a “cytokine storm”. While antimicrobials such as antibiotics and antiviral agents effectively reduce pathogen burden during infections, once hyperinflammation is initiated, the risk of progression to sepsis remains irrespective of pathogen clearance. This highlights the need for targeted interventions to manage inflammatory responses alongside pathogen eradication. Conventional immunosuppressive therapies, like glucocorticoids that are effective in many inflammatory conditions, fail to consistently improve outcomes in septic patients^1,2^. Similarly, biologic therapies targeting individual cytokines such as TNFα or IL-6 have shown limited success, underscoring the multifactorial nature of sepsis and the inadequacy of targeting discrete inflammatory mediators^3^. Thus, a deeper understanding of the pathways governing immune activation is critical for the development of novel therapeutics to dampen systemic inflammation in sepsis.

Recent advances have highlighted that immune cell metabolism extends beyond energy production and biosynthesis. Dynamic changes in metabolites and their intermediates shape cellular function and fate, influencing both the activation state and effector functions of immune cells. For instance, enhanced glycolytic flux in activated CD8^+^ T cells channels metabolites into the pentose phosphate pathway, fueling NADPH required for reduced glutathione production and subsequent ROS control^4^. Beyond serving as metabolic substrates, certain metabolites act as signaling molecules, and excessive accumulation can induce post-translational modifications of proteins, thereby modulating immune responses^5–7^. Although the Warburg effect, where cells preferentially utilize glycolysis despite the availability of oxygen, is well documented in macrophage activation, the roles of alternative metabolic pathways are still not well characterized. The metabolic characterization has also been performed mainly on mouse bone marrow-derived macrophages (BMDMs) in vitro, but the immunometabolic mechanisms in monocytes in vivo are not well understood.

The electron transport chain (ETC) is the central machinery for adenosine triphosphate (ATP) production. It comprises four complexes (complex I-IV) that transfer electrons down the chain to generate a proton gradient that is utilized by ATP synthase to produce ATP. Immunological functions have increasingly been ascribed to the ETC, as alterations in its activity can shape immune cell behaviour^8,9^. For example, suppression of the ETC by nitric oxide and diversion of the metabolites in the Tricarboxylic Acid (TCA) cycle helps to switch the cells from aerobic respiration to glycolysis during macrophage activation^10,11^. Additionally, reactive oxygen species (ROS) generated by complexes I and III activates inflammatory pathways such as NF-κB and HIF1α^12,13^. We and others have also shown that perturbing the ETC complexes attenuates cytokine secretion after inflammatory trigger^9,14^. Deletion of individual complexes also exerts distinct effects on leukocyte subsets, highlighting the nuanced role of each ETC component in immune regulation^15,16^.

We and others previously identified MOCCI (Modulator of Cytochrome C Oxidase during Inflammation), a subunit of CIV encoded by the gene *C15orf48* ^9,17,18^. We showed that MOCCI expression is selectively induced upon IL-1β stimulation in human endothelial cells (ECs) and lung epithelial cell lines. Interestingly, *C15orf48* also harbours the microRNA hsa-mir-147b. Following inflammatory activation, both MOCCI and miR-147b are upregulated. MOCCI replaces its paralog NDUFA4 (NADH dehydrogenase [ubiquinone] 1 alpha subcomplex subunit 4) in CIV, while miR-147b promotes degradation of *NDUFA4* transcripts, collectively resulting in reduced CIV activity in these cell types. Functionally, MOCCI and miR-147b exert anti-inflammatory effects in endothelial and epithelial cells by limiting cytokine secretion, although they have opposing impacts on cell death and the interferon response, suggesting that miR-147b has other targets beyond *NDUFA4*. The combination of MOCCI and miR-147b strikes a balance between rapid pathogen clearance while ensuring cell survival in endothelial and epithelial cells. Several mouse studies have since shown that *C15orf48* is needed to suppress inflammation in the colon. Knocking it out exacerbates DSS-induced colitis and regeneration after colon injury^19,20^. *C15orf48* has also been implicated in tumour immunity, where it is upregulated in tumour-associated macrophages and knocking it out augmented tumour growth^21^. Beyond regulating the cellular immune response, *C15orf48* is also required for stress-independent autophagy in thymic epithelial cell function and knocking it out impairs self-tolerance, leading to an autoimmune phenotype^22^.

Here we report that upon challenge with *Klebsiella pneumoniae* (Kp) or lipopolysaccharide (LPS), *C15orf48-*knockout (KO) mice had lower systemic inflammation and higher survival rates. Contrary to what we observed in endothelial cells, deletion of *C15orf48* in myeloid cells attenuates systemic inflammation by suppressing monocyte activation. Endothelial-specific deletion of *C15orf48* results in exacerbated inflammation, indicating that *C15orf48* plays cell type-specific functions. Metabolomic analyses in monocytes revealed that *C15orf48* is essential for regulating lipid and amino acid metabolism, uncovering a Complex IV (CIV)– fatty acid oxidation–branched-chain amino acid link that underpins robust monocyte activation. These findings provide new insights into the metabolic control of inflammation and identify *C15orf48* as a potential therapeutic target for modulating monocyte-driven immune responses in sepsis.

## Results

### C15orf48-KO mice are more resistant and tolerant to sublethal Klebsiella infection

To assess whether *C15orf48* is upregulated in vivo during bacteria-induced inflammation, we induced systemic inflammation in mice through intraperitoneal injection of LPS. *C15orf48* expression was increased in all tissues examined except the colon, which displayed constitutively high baseline expression (Fig. 1a). Among these, the lung exhibited the greatest induction of *C15orf48* (Fig. 1a), prompting us to further investigate the functional role of *C15orf48* in respiratory infection models.

**Figure 1.**
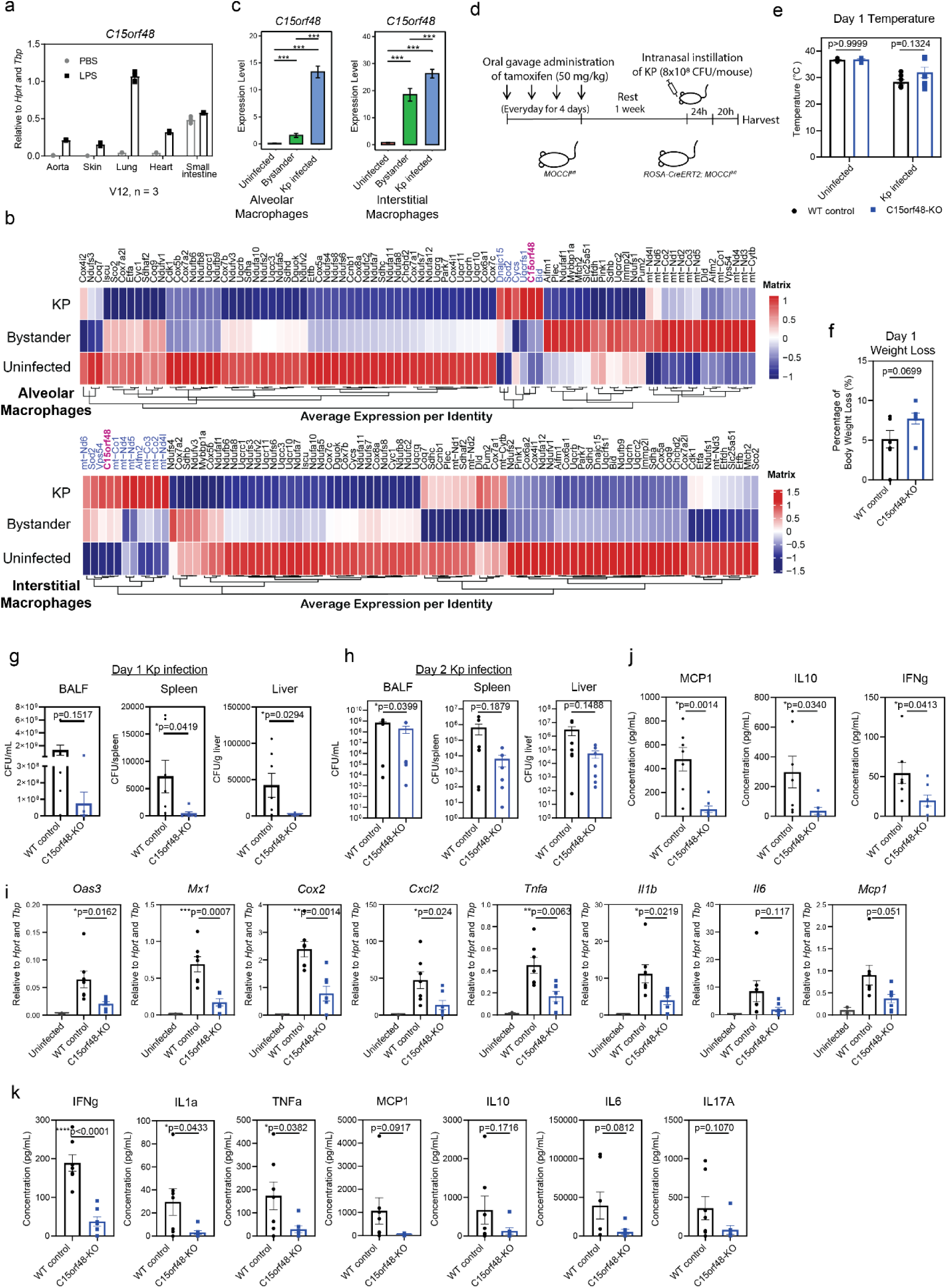
*C15orf48-*KO mice are less susceptible to Kp infection. (a) qPCR of *C15orf48* in aorta, skin, lung, heart and small intestine of mice after PBS and LPS injection. (n=3) (b) Expression of ETC genes, based on scRNA-seq of alveolar and interstitial macrophages isolated from mouse lungs. These mice were either uninfected (Uninfected) or infected with Kp. The cells from infected mice were further annotated as those with Kp identified within them (Kp infected) and those without Kp (Bystander). *C15orf48* is highlighted in purple, while upregulated genes in infected cells are highlighted in blue. Data was taken from GSE184290^23^. (c) *C15orf48* expression in alveolar and interstitial macrophages in uninfected mice (Uninfected) and Kp infected mice (Kp-infected, Bystander). Data was taken from GSE184290^23^. (d) Schematic showing the administration of tamoxifen and intranasal infection of WT and *C15orf48-*KO mice with Kp. (e) Body temperature of WT and *C15orf48-*KO mice before and one day after Kp infection. P = two-way ANOVA; n = 7 biological replicates over two repeats. (f) Percentage weight loss of WT and *C15orf48-*KO mice one day after Kp infection. P = two-tailed unpaired Student’s t test; n = 7 biological replicates over two repeats. (g) Quantification of colony forming units (CFU) in BALF, spleen and liver from WT and *C15orf48-*KO mice one day after Kp infection. Uninfected mice had no CFU and are not shown. P = two-tailed unpaired Student’s t test; n = 7 biological replicates over two repeats. (h) Quantification of colony forming units (CFU) in BALF, spleen and liver from WT and *C15orf48-*KO mice two days after Kp infection. Uninfected mice had no CFU and are not shown. P = two-tailed unpaired Student’s t test; n = 8 biological replicates over two repeats. (i) qPCR of lungs from WT or *C15orf48-*KO mice one day after Kp infection. P = two-tailed unpaired Student’s t test between WT and *C15orf48-*KO mice; n = 7 biological replicates over two repeats. (j) Cytokine analysis of BALF taken from WT or *C15orf48-*KO mice one day after Kp infection. P = two-tailed unpaired Student’s t test; n = 7 biological replicates over two repeats. (k) Cytokine analysis of plasma taken from WT or *C15orf48-*KO mice one day after Kp infection. P = two-tailed unpaired Student’s t test; n = 7 biological replicates over two repeats.

We selected *Klebsiella pneumoniae* (Kp), a major cause of hospital-acquired pneumonia and well-documented for its antibiotic resistance, as a clinically relevant pathogen. Single-cell RNA sequencing (scRNA-seq) of mouse lungs during Kp infection (GSE184290^23^) demonstrated that infected alveolar and interstitial macrophages downregulated ETC genes (Fig. 1b). Notably, a subset of ETC genes including *C15orf48*, was upregulated specifically in infected macrophages, and much lower in bystander populations (Fig. 1b,c), representing cell-autonomous responses to infection. In our previous studies in human ECs, *C15orf48* induction led to reduced CIV activity, which is congruent with the observed general downregulation of ETC genes seen here.

To investigate the effect of *C15orf48* induction during infection, we generated inducible *C15orf48*-KO mice by crossing ROSA-creERT2 with *C15orf48*-floxed lines^24,25^. LoxP sites were inserted upstream of exon 2 and downstream of exon 4. Tamoxifen administration resulted in KO of both MOCCI and miR-147b in all organs tested (Suppl. Fig. 1a–c). Wild-type (WT) and *C15orf48-*KO mice were challenged with 8 × 10^8^ CFU Kp (MGH78578), a dose that elicits transient illness (days 1–2) followed by recovery (days 3–4)(Fig. 1d). No significant differences in body temperature or weight loss were observed between genotypes, although KO mice exhibited a trend toward slightly higher body temperature and weight loss at 1 day post infection (dpi) that normalized by 2 dpi (Fig. 1e,f; Suppl. Fig. 1d,e).

Remarkably, *C15orf48-*KO mice exhibited a significantly lower bacterial burden: reduced colony counts were detected in liver and spleen on 1 dpi, and in bronchoalveolar lavage fluid (BALF) on 2 dpi post-infection (Fig. 1g,h). These findings correlated with decreased expression of proinflammatory genes in KO lungs at 1 dpi (Fig. 1i).

Given the attenuated cytokine transcript levels in KO lungs, we assayed the BALF at 1 dpi for 13 inflammatory mediators. MCP1, IFNγ, and IL-10 were significantly reduced, while TNFα, IL-1β, IL-6, IL-1α, IL-27, and IL-12p70 showed a similar downward trend (Fig. 1j; Suppl. Fig. 1f). Reduced local inflammation was mirrored systemically, as circulating IFNγ, IL-1α, and TNFα were also diminished in KO mice (Fig. 1k). By 2 dpi, these differences between genotypes were less pronounced, indicating a predominant role for *C15orf48* during early infection (Suppl. Fig. 1g-i).

### *C15orf48* is required for a robust pro-inflammatory response during endotoxemia

A reduction in local and systemic cytokine production could be a consequence of enhanced bacterial clearance in earlier stages of infection. However, since we measured both bacterial burden and cytokine production in *C15orf48-*KO mice within 18 h of infection and hence very early during infection (Day 1), the observed concurrent dual reduction is paradoxical, since enhanced bacterial clearance is usually supported by a more robust immune response^26^. To delineate whether the reduced systemic inflammation observed in *C15orf48-*KO mice is due to suppressed immune response or lower bacterial load (i.e, to isolate the acute inflammatory response independent of infection), we induced an acute innate response in WT and *C15orf48-*KO mice via intraperitoneal injection of LD50 (Lethal Dose 50%) dose of LPS (Fig. 2a), an endotoxin derived from the outer membrane of gram negative bacteria that induces lethal hypercytokinemia in mice and is an established model for sepsis.

**Figure 2.**
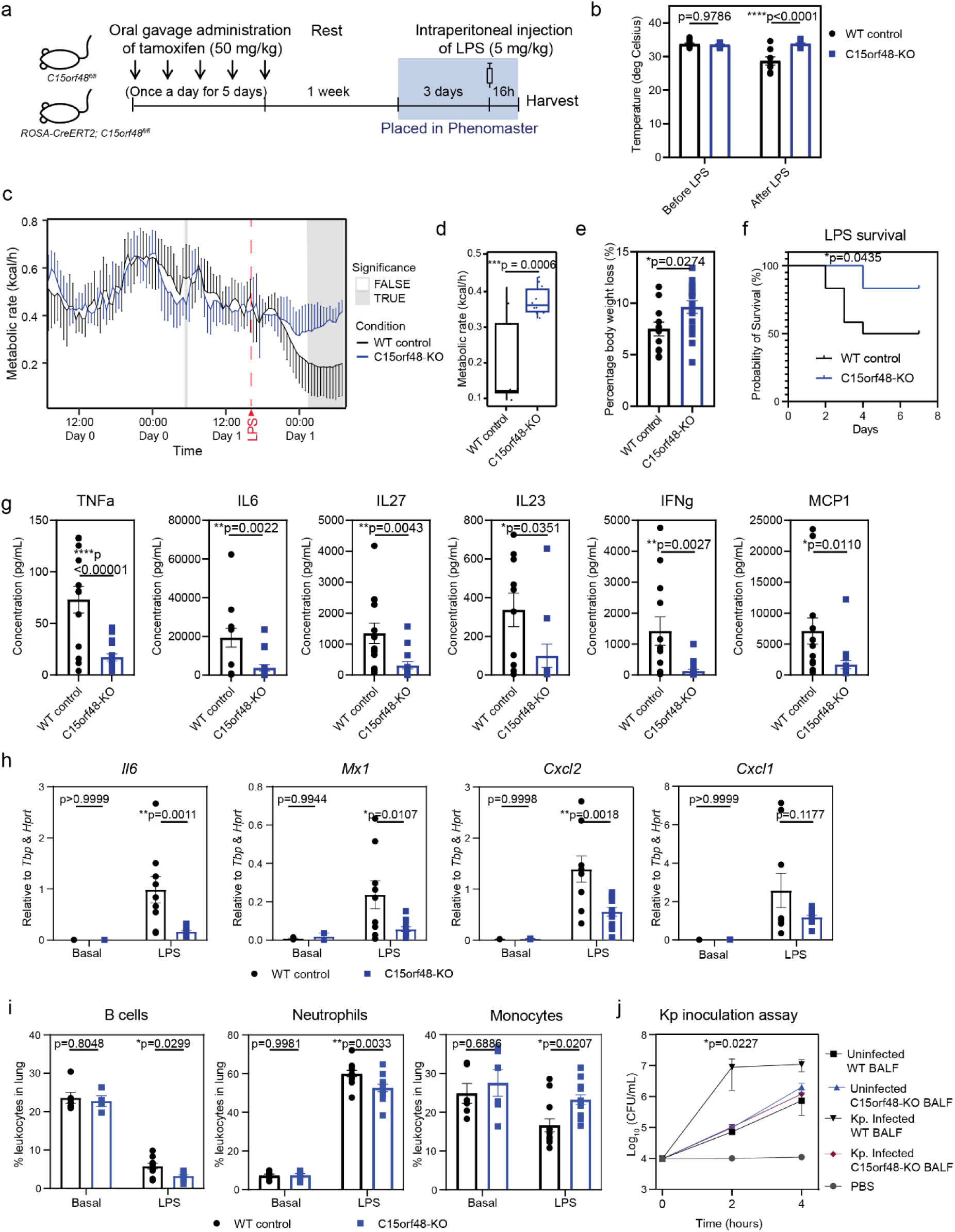
*C15orf48-*KO mice exhibit lower systemic inflammation during endotoxemia. (a) Schematic of WT and *C15orf48-*KO mice treated with tamoxifen, before they were placed in the PhenoMasterand administered with LPS. (b) Body temperature of WT and *C15orf48-*KO mice before and 16 hours after LPS injection. P = two-way ANOVA; n = 12 biological replicates over two repeats. (c) Metabolic rate of WT and *C15orf48-*KO mice before and after LPS infection, as measured by the PhenoMaster. P = two-way ANCOVA; n = 10 biological replicates over two repeats. (d) Median metabolic rate of WT and *C15orf48-*KO mice 10 hours after LPS infection, as measured by the PhenoMaster. P = two-way ANCOVA; n = 10 biological replicates over two repeats. (e) Percentage weight loss of WT and *C15orf48-*KO mice 16 hours after LPS injection. P = two-tailed unpaired Student’s t test; n = 16 biological replicates over three repeats. (f) Survival curve of LPS-injected mice over 7 days. P = Gehan-Breslow-Wilcoxon test; n = 12 biological replicates over two repeats. (g) Cytokine analysis of plasma taken from WT or *C15orf48-*KO mice 16 hours after LPS injection. P = two-tailed unpaired Student’s t test; n = 12 biological replicates over two repeats. (h) qPCR of lungs from WT or *C15orf48-*KO mice 16 hours after LPS injection. P = two-way ANOVA; n = 4 and 8 biological replicates for basal and after LPS, respectively, over two repeats. (i) Flow cytometry analysis of infiltrated leukocytes in lungs from mice injected with PBS or LPS. P = two-way ANOVA; n = 6 and 12 biological replicates for basal and after LPS, respectively, over three repeats. (j) Quantification of Kp growth in bacterial-cleared BALF from uninfected and previously-infected mice. P = two-way ANOVA; n = 7 biological replicates per condition from two repeats.

After LPS injection, WT mice gradually reduced their metabolism, resulting in a lower body temperature measured 16h after the injection (Fig. 2b,c). Remarkably, the *C15orf48-*KO mice recovered from the hypometabolism 7 hours after the LPS injection (Fig. 2b-d). As a result, the knockout mice were able to re-establish homeothermy (Fig. 2b) without significant differences in feeding or activity (Suppl. Fig. 2a,b). MOCCI is not expressed in the skeletal muscle, ruling out the direct effect of MOCCI on muscle metabolism or shivering thermogenesis (Suppl. Fig. 2c). The remarkable ability of the KO mice to maintain temperature despite isocaloric intake did have trade-offs, as they exhibited greater weight loss (Fig. 2e). KO mice were also visibly less lethargic and more responsive (Supplementary Video 1), which we interpret as a lower sickness score. Consequently, *C15orf48-*KO mice had a significantly lower mortality rate compared to WT mice, suggesting that they are protected against LPS-induced lethal sepsis (Fig. 2f).

As sepsis is accompanied by hypercytokinemia, we compared the cytokine profile between the WT and KO mice. Cytokine levels in the plasma (Fig. 2g) and transcripts of pro-inflammatory genes in lungs were lower in the KO mice (Fig. 2h). This was accompanied by fewer neutrophils and B cells recruited to the lungs, although monocyte recruitment was elevated (Fig. 2i). Curiously, there are fewer T cells in *C15orf48-*KO lungs at basal but these differences were eliminated after LPS (Suppl. Fig. 2d). As Takakura et al. has shown that *C15orf48* plays a role in thymic epithelial cells, the reduction in T cells could possibly be due to developmental defects in the thymus^27^. As T cells are recruited later in an infection, they are unlikely to be responsible for the suppressed acute immune response. Overall, these results demonstrate that *C15orf48*–KO mice possess an intrinsically less pro–inflammatory innate immune response, independent of bacterial load, as shown by their reduced systemic and local inflammation following LPS challenge. However, this draws the question of how KO mice have better bacterial control (Fig. 1g,h) despite a reduced innate inflammatory response.

Previous work in acute respiratory distress syndrome (ARDS) has shown that intrapulmonary cytokines released during infection can directly enhance the growth of clinically relevant bacteria in nosocomial infections^28,29^. In the lung, Kp siderophores have been shown to directly induce IL–6, CXCL1, and CXCL2 secretion which in turn drive bacterial dissemination to the spleen via HIF–1α stabilization, independently of bacterial burden^30^. Similar principles have emerged from metabolomic studies, where host–derived metabolites can benefit bacterial fitness^31^, illustrating how opportunistic pathogens exploit host inflammatory and metabolomic signals to their advantage. Altogether, this points to a bidirectional loop between host and pathogen in which dysregulated inflammation favors reproduction of some opportunistic bacteria.

To address if this is the reason why *C15orf48*–KO mice have a lower Kp burden despite reduced lung inflammation, we examined how Kp grows following inoculation into BALF harvested from WT and *C15orf48*–KO mice. BALF from healthy WT and *C15orf48-*KO mice supported equally enhanced Kp growth over PBS alone, indicating that BALF contains factors that promote Kp replication (Fig. 2j). Strikingly, Kp inoculated into BALF from previously infected WT mice grew two log-fold faster than Kp inoculated into BALF from uninfected mice, indicating that Kp infection induces a lung milieu that supports its own growth. It is important to note that BALF collected from previously infected mice was cleared of bacteria during collection before inoculation, since parallel BALF cultures without secondary Kp inoculation did not show bacterial growth, ensuring that subsequent growth reflected the newly inoculated bacteria rather than carryover from the original infection. In contrast, Kp growth in BALF from previously infected *C15orf48*–KO mice was significantly lower, resembling growth in BALF from uninfected animals, suggesting that the KO lung environment fails to acquire a pro–bacterial state (Fig. 2j), explaining the reduced Kp load observed in *C15orf48*–KO mice.

Although identifying the specific soluble factors in BALF that renders the WT milieu more pro–bacterial than KO mice would be of considerable interest, dissecting these components is beyond the scope of this study. Instead, we focus on understanding why *C15orf48* deficiency makes the mice intrinsically less pro–inflammatory.

### C15orf48 is upregulated in myeloid and ECs during endotoxemia

To determine the cellular mechanisms underlying the suppressed immune response in the lungs, we performed scRNA-seq on lung tissue from WT and *C15orf48-*KO mice following intraperitoneal injection with either PBS or LPS (Fig. 3a) and delineated 9 cell types according to cell-type specific markers in both conditions (Suppl. Fig. 3a). scRNA-seq analysis revealed that *C15orf48* expression is specifically induced after LPS treatment and is restricted to distinct cell populations, namely monocytes, neutrophils, and ECs (Fig. 3b,c). As expected, cells derived from KO mice exhibited minimal to absent *C15orf48* expression, confirming the efficacy of the knockout strategy (Suppl. Fig. 3b).

**Figure 3.**
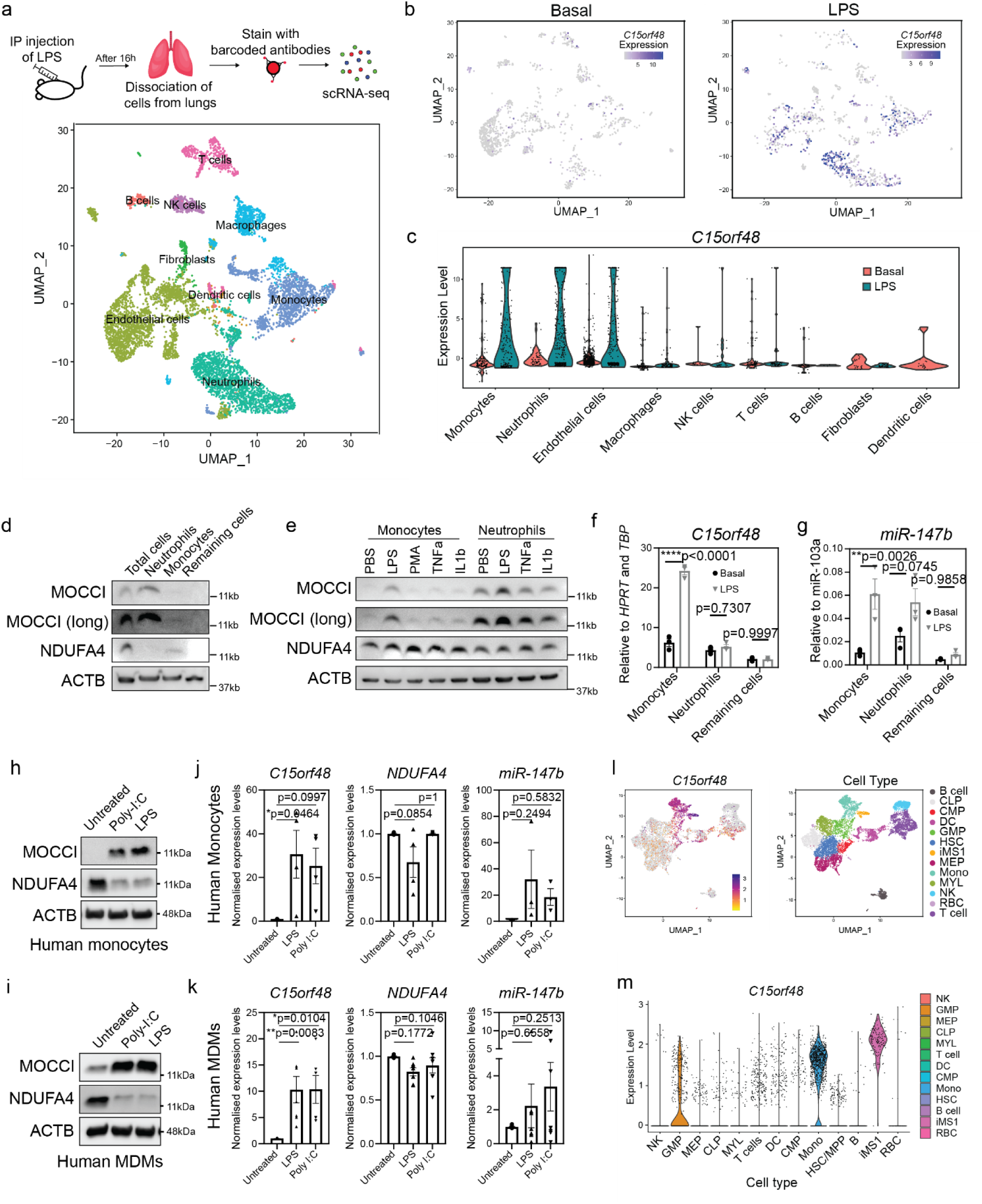
*C15orf48* is expressed in human and mouse monocytes. (a) (Top) Schematic of the process to obtain single cell isolates from lungs. (Bottom) UMAP of the cell types identified from the scRNA-seq done on the single cell isolates from lungs. (b) Feature plot of *C15orf48* expression in single cell lung isolates from mice injected with PBS and LPS. *C15orf48* is annotated as AA467197 in mice. (c) Violin plot of *C15orf48* expression in single cell lung isolates from mice injected with LPS. (d) Western blot of MOCCI and NDUFA4 in different cells isolated from mouse bone marrow. “Remaining cells” are remaining bone marrow cells after neutrophil and monocyte depletion. n = 4 biological repeats. (e) Western blot of MOCCI and NDUFA4 in murine neutrophils and monocytes after 16 hours treatment with LPS, PMA, TNFa and IL1b. n = 3 biological repeats. (f) qPCR of *C15orf48* in monocytes, neutrophils and bone marrow cells depleted of monocytes and neutrophils (“remaining cells”) after 16h treatment with LPS or vehicle. P = two-way ANOVA; n = 3 biological repeats. (g) qPCR of miR-147b in monocytes, neutrophils and bone marrow cells depleted of monocytes and neutrophils (“remaining cells”) after 16h treatment with LPS or vehicle. P = two-way ANOVA; n = 3 biological repeats. (h) Western blot of MOCCI and NDUFA4 in human monocytes after 16 hours treatment with LPS and poly-I:C. n = 4 biological repeats. (i) Western blot of MOCCI and NDUFA4 in human monocyte-derived-macrophages after 16 hours treatment with LPS or poly-I:C. n = 6 biological repeats. (j) qPCR of *C15orf48, NDUFA4* and miR-147b in human monocytes after 16h treatment with LPS or poly-I:C. P = one-way ANOVA; n = 4 biological repeats. (k) qPCR of *C15orf48, NDUFA4* and miR-147b in human monocyte-derived-macrophages after 16h treatment with LPS or poly-I:C. P = one-way ANOVA; n = 6 biological repeats. (l) (l-m) UMAP (l) and violin plot (m) showing expression of *C15orf48* in human bone marrow mononuclear cells from septic patients. Data was taken from SCP550 from Broad Institute’s Single Cell Portal^32^. Cell legend: iMS1 = Inflammatory Monocyte State 1; B = B cells; DC = Dendritic cells; Mono = Monocytes; NK = Natural Killer cells; RBC = Red blood cells; T cells = T cells; CLP = Common Lymphoid Progenitor; CMP = Common Myeloid Progenitor; GMP = Granulocyte-Macrophage Progenitor; HSC = Hematopoietic Stem Cell; MEP = Megakaryocyte-Erythroid Progenitor.

As we had previously shown the expression of MOCCI and miR-147b in human ECs, we focused on confirming their expression in neutrophils and monocytes, using myeloid cells from mouse bone marrow and human blood as a proxy for lung-infiltrating myeloid cells, since the latter are derived from circulating cells^9^. Freshly isolated neutrophils from bone marrow show strong expression of MOCCI, while monocytes had relatively weaker expression and instead express NDUFA4, the paralog of MOCCI (Fig. 3d). After 16h culture in LPS, both cell types upregulate MOCCI, with a larger relative increase in monocytes compared with basal (Fig. 3e). This pattern was mirrored in the expression of *C15orf48* transcripts and miR-147b, where they are upregulated strongly in monocytes and not as drastically in neutrophils (Fig. 3f,g). Mouse BMDMs exhibited high levels of MOCCI that was eliminated by KO of *C15orf48*, which further increased upon LPS stimulation in WT (Suppl. Fig. 3c). Notably, both human PBMC monocytes and monocyte-derived macrophages upregulated *C15orf48* mRNA and MOCCI protein in response to LPS and poly(I:C) stimulation (Fig. 3h-k), indicating a conserved induction of *C15orf48* in the myeloid compartment during an infectious challenge. This was accompanied by a reduction in NDUFA4 protein levels, coherent with our previous findings in endothelial and epithelial cells that the levels of these two proteins are negatively correlated. The reduction in NDUFA4 cannot be accounted for by the modest reduction in NDUFA4 transcripts, indicating that NDUFA4 protein level is regulated post-transcriptionally (Fig. 3j,k).

Similar to what we observed in the mouse, scRNA-seq of human bone marrow-derived monocytes treated with LPS also showed high *C15orf48* expression in a subset of monocytes (iMS1) that are also in septic patients (Fig. 3l,m)^32^. Other groups have also reported high expression of *C15orf48* in monocytes in rheumatoid arthritis and COVID-19 patients^33^. These findings provide evidence that *C15orf48* is upregulated in human monocytes in several inflammatory disease states.

In summary, *C15orf48* and MOCCI are upregulated specifically and drastically in ECs and monocytes following infectious challenge in both mouse (lung, bone marrow) and human (bone marrow, blood).

### Knockout of *C15orf48* in myeloid cells but not ECs reduce systemic inflammation during endotoxemia

To decipher the relative contribution of *C15orf48* in ECs, monocytes and neutrophils to LPS-induced sepsis, we generated cell-specific KO mouse models. For myeloid deletion, we generated *C15orf48-*fl/fl; LysM-Cre mice (*C15orf48*^m^-KO); for ECs, we generated *C15orf48-*fl/fl; Cdh5-creERT2 mice (*C15orf48*^EC^-KO)^34^.

Surprisingly, *C15orf48*^EC^-KO led to significant increase in proinflammatory gene expression and reduced weight loss following LPS challenge (Fig. 4a,b), in opposing contrast to the global *C15orf48-*KO phenotype (Fig. 2b,e). While circulating pro-inflammatory cytokines were slightly elevated in these mice, the differences did not reach statistical significance (Suppl. Fig. 4a). These findings indicate that *C15orf48* exerts an anti-inflammatory effect in ECs, consistent with its role we previously reported in primary human aortic ECs^9^.

**Figure 4.**
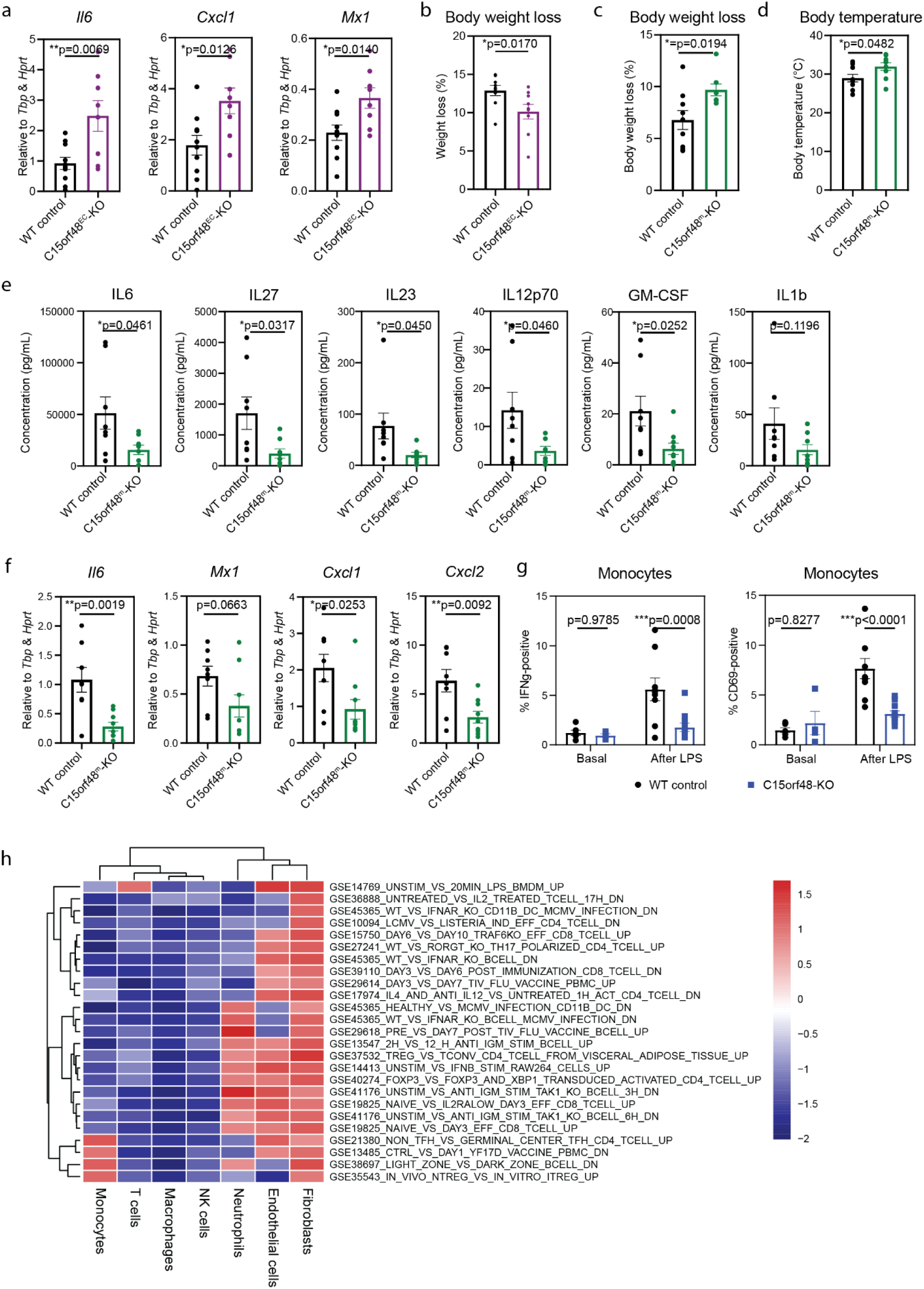
*C15orf48* plays contradictory roles in myeloid and endothelial cells during LPS-induced systemic inflammation. (a) qPCR of pro-inflammatory genes in lungs from WT and *C15orf48*^EC^-KO mice after LPS injection. P = two-tailed unpaired Student’s t test; n = 9 biological replicates over two repeats. (b) Percentage weight loss of WT and *C15orf48*^EC^-KO mice one day after LPS injection. P = two-tailed unpaired Student’s t test; n = 9 biological replicates over two repeats. (c) Percentage weight loss of WT and *C15orf48*^m^-KO mice one day after LPS injection. P= two-tailed unpaired Student’s t test; n = 8 biological replicates over two repeats. (d) Temperature of WT and *C15orf48*^m^-KO mice one day after LPS injection. P = two-tailed unpaired Student’s t test; n = 8 biological replicates over two repeats. (e) Levels of proinflammatory cytokines in plasma of WT and *C15orf48*^m^-KO mice one day after LPS injection. P = two-tailed unpaired Student’s t test; n = 8 biological replicates over two repeats. (f) qPCR of pro-inflammatory genes in lungs from WT and *C15orf48*^m^-KO mice after LPS injection. P = two-tailed unpaired Student’s t test; n = 8 biological replicates over two repeats. (g) Percentage of IFNγ- and CD69-positive monocytes in lungs from WT and *C15orf48-*KO mice at basal and after LPS injection. P = two-way ANOVA; n = 6 biological replicates for mice at basal and 9 for LPS-injected mice from three repeats. (h) GSEA analysis of *C15orf48-*KO versus WT in different cell types, based on scRNA-seq of cells isolated from lungs after LPS injection.

Conversely, when *C15orf48* was knocked out in the myeloid cells, *C15orf48*^m^-KO mice phenocopied the global KO mouse, with a higher body temperature and higher weight loss after LPS injection compared to control (Fig. 4c,d); lower levels of pro-inflammatory plasma cytokine (Fig. 4e; Suppl. Fig. 4b) and pro-inflammatory genes in their lungs (Fig. 4f). Notably, the same genes that are lower in whole body *C15orf48-*KO mice are also lower in *C15orf48*^m^-KO (Fig. 2g,h vs Fig. 4e,f), indicating that the lower systemic inflammation in response to LPS observed in global *C15orf48-*KO mice is mainly driven by the KO of *C15orf48* in myeloid cells. To determine if myeloid cells are less activated in *C15orf48-*KO mice, we measured the levels of IFNg and CD69 on lung isolates from LPS/PBS-injected mice via flow cytometry. Despite the increased number of monocytes in the global *C15orf48-*KO lungs (Fig. 2i), these monocytes exhibited reduced activation relative to WT controls (Fig. 4g). This was not observed in neutrophils, where IFNg was equally induced in both WT and KO cells (Suppl. Fig. 4c). Although *C15orf48*^EC^-KO mice had higher systemic inflammation, flow analysis of ECs from global *C15orf48-*KO mice – where both monocytes and EC are *C15orf48* deficient - nonetheless revealed reduced EC activation after LPS treatment, as evidenced by lower VCAM1 expression (Suppl. Fig. 4d). This implies that the pro-inflammatory role of *C15orf48* in monocytes is cell-autonomous and exerts a dominant role over its anti-inflammatory effect in non-immune cell types like ECs.

Surprised by the potential divergent functions of *C15orf48* in different cell types, we performed gene set enrichment analysis (GSEA) of immunological pathways on the scRNA-seq of the WT and *C15orf48-*KO lungs (Fig. 4h). *C15orf48-*KO monocytes and macrophages clearly display the opposite trend from ECs, where they exhibited a less and more inflammatory gene signature, respectively. *C15orf48-*KO neutrophils showed a more nuanced profile, with positive associations in certain inflammatory pathways. This suggests that *C15orf48* might play different roles even within the myeloid compartment. To further investigate the cell-specific functions of *C15orf48*, we performed GSEA on bulk RNA-seq data from organs with high *C15orf48* expression. We correlated expression of *C15orf48* with different gene signatures in each organ. Notably, tissues where *C15orf48* is predominantly expressed in leukocytes (whole blood) has opposite trend compared to tissues where the expression is in non-leukocytes (testes, stomach)(Suppl. Fig. 4e), underscoring the fact that *C15orf48* function is context- and cell-type-specific.

### *C15orf48* is required for robust monocyte activation

We first assessed the bioenergetics of WT and KO myeloid cells. To our surprise, there were no detectable differences in CIV activity between freshly isolated bone marrow WT and KO neutrophils and monocytes (Fig. 5a,b) or BMDMs (Suppl. Fig. 5a), contrary to our previous data in human ECs where MOCCI overexpression led to reduced CIV activity. One possible explanation is that the sensitivity of the assay is insufficient to detect differences in cells with very low CIV activity such as these or the presence of compensatory mechanisms in these myeloid populations. Despite the lack of differences in CIV activity, *ex vivo* untreated *C15orf48-*KO monocytes had reduced basal and maximal respiration (Fig. 5c-e). This difference was normalized following LPS treatment, which suppresses OCR in monocytes, similar to what happens in BMDMs during LPS treatment (Fig. 5c-e)^35^. Conversely, while untreated monocytes had no differences in glycolytic rate (Fig. 5f), KO monocytes had a lower glycolytic rate following LPS treatment (Fig. 5g). This is consistent with the findings that they are less activated, as evidenced by lower levels of IL1b, IL6 and GM-CSF in response to LPS than WT monocytes (Fig. 5h). Altogether, the data suggests that the KO of *C15orf48* in monocytes affects its intrinsic respiration which leads to reduced activation during LPS challenge. Despite high levels of MOCCI expression in neutrophils, *C15orf48-*KO neutrophils did not show any difference in their respiration, glycolytic rate and oxidative burst, nor in their IFNg levels (Suppl. Fig. 4c, 5b-d). As such, going forward, we focused on monocytes.

**Figure 5.**
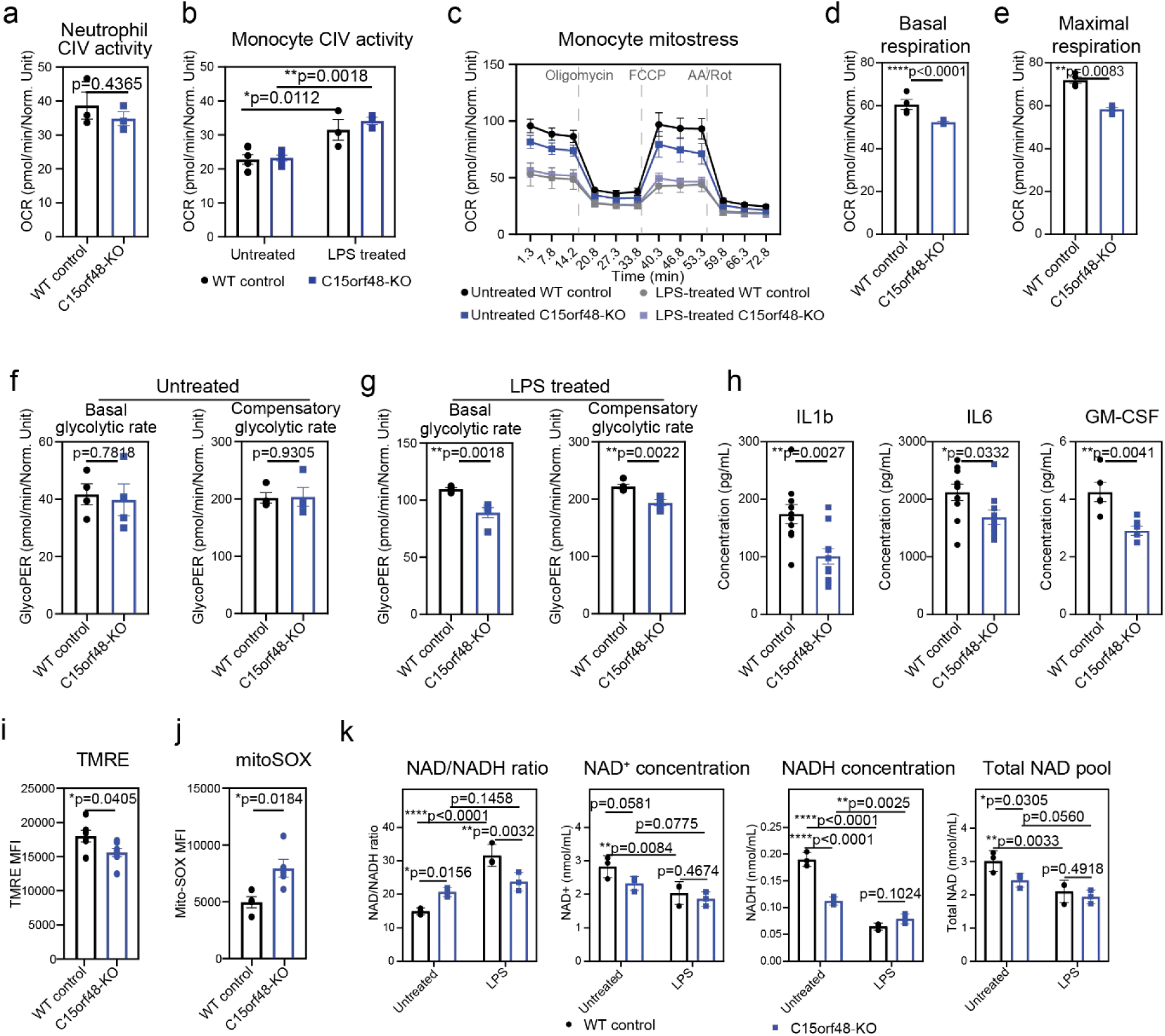
*C15orf48* deficiency alters bioenergetics, redox balance, and cytokine secretion in monocytes. (a) Complex IV activity of WT and *C15orf48-*KO murine neutrophils at basal. P = two-tailed unpaired Student’s t test; n = 3 biological replicates. (b) Complex IV activity of WT and *C15orf48-*KO murine monocytes at basal and after 6h LPS treatment. P = two-way ANOVA; n = 6 biological replicates from two repeats at basal and 3 biological replicates per condition after LPS treatment. Each biological replicate contained cells pooled from 2-3 mice. (c) Seahorse XF Cell Mito Stress Test assay on WT and *C15orf48-*KO monocytes at basal and after 6h LPS treatment. n = 4 biological replicates per condition from two repeats. Each biological replicate contained cells pooled from 2-3 mice. (d-e) Basal respiration (d) and maximal respiration (e) of WT and *C15orf48-*KO monocytes after 6h LPS treatment, as obtained from the Mito Stress assay in (c). P = two-tailed unpaired Student’s t test; n = 4 biological replicates over two repeats. Each biological replicate contained cells pooled from 2-3 mice. (f-g) Basal and compensatory glycolytic rate of WT and *C15orf48-*KO monocytes at basal (f) and after 6h LPS treatment (g), as measured by Seahorse XF Glycolytic Rate assay. P = two-tailed unpaired Student’s t test; n = 4 biological replicates over two repeats. Each biological replicate contained cells pooled from 2-3 mice. (h) Cytokine secretion by WT and *C15orf48-*KO monocytes after 6h LPS treatment. P = two-tailed unpaired Student’s t test; n = 10 biological replicates over two repeats for IL1b and IL6. n = 4 biological replicates for GM-CSF. (i) Quantification of mitochondrial membrane potential in WT and *C15orf48-*KO monocytes at basal, based on TMRE dye staining. P = two-tailed unpaired Student’s t test; n = 6 biological replicates over two repeats. (j) Quantification of mitochondrial ROS levels in WT and *C15orf48-*KO monocytes at basal, based on mitoSOX™ dye staining. P = two-tailed unpaired Student’s t test; n = 5 biological replicates over two repeats. (k) Quantification of NAD/NADH ratio, NAD^+^, NADH and total NAD concentrations in WT and *C15orf48-*KO monocytes at basal and 6h after LPS treatment. P = two-way ANOVA; n = 3 biological replicates.

Since *C15orf48-*KO monocytes have clear differences in their metabolism compared to WT, we next sought to define the mechanistic relationship between the altered metabolism and dampened activation. Coherent with their reduced respiration (Fig. 5d,e), *C15orf48-*KO monocytes had lower mitochondrial membrane potential (Fig. 5i) that is accompanied by increased mitochondrial ROS (Fig. 5j), indicating an overall reduction in the efficiency of the ETC in *C15orf48-*KO monocytes that might be the cause of dampened activation. As ETC efficiency is tightly coupled to the cellular NAD/NADH redox state, and myeloid activation requires adequate replenishment of the NAD pool^36^, we hypothesized that NAD availability and/or redox balance might be altered in *C15orf48*–KO monocytes. We therefore quantified NAD^+^ and NADH levels in WT and *C15orf48*–KO cells at baseline and after LPS stimulation . Consistent with previous reports^36^, LPS reduced both NAD^+^ and NADH in WT and KO monocytes, but only WT cells displayed a marked increase in the NAD^+^/NADH ratio (Fig. 5k). In contrast, KO monocytes exhibited a smaller total NAD pool and a more oxidized NAD^+^/NADH ratio already at baseline, which is consistent with their reduced respiratory capacity (Fig. 5d,e,k). Notably, both the size of the NAD pool and the NAD^+^/NADH ratio changed less dynamically in KO monocytes than in WT cells following LPS, indicating a diminished capacity to remodel redox cofactors during activation (Fig. 5k). While the muted NAD changes in KO after LPS stimulation could be a reflection of weaker activation, rather than defects in regulating NAD metabolism, the smaller NAD pool at basal is more supportive of the latter scenario. Given that reduced NAD availability impairs GAPDH activity, a key rate–limiting glycolytic enzyme required for macrophage activation^36,37^, these findings align with the attenuated glycolytic response observed in KO monocytes after LPS stimulation.

In summary, loss of *C15orf48* leads to reduced respiration, increased mitochondrial ROS, and a smaller, more oxidized NAD pool at baseline, leaving KO monocytes with limited capacity to reconfigure NAD/NADH during LPS–induced activation. This constrained redox flexibility is expected to impair multiple NAD–dependent catabolic pathways such as lipid and amino–acid metabolism, which might in turn contribute to the attenuated inflammatory response of KO monocytes.

### *C15orf48* is required for fatty acid oxidation in monocytes

To identify which downstream pathways are affected, we next performed targeted metabolomics on WT and *C15orf48*–KO monocytes that were either untreated or treated with LPS for 4 hours. Out of the 326 targeted metabolites, 102 were detected (Suppl. Table 1). Principle component analysis (PCA) of the detected metabolites revealed that WT and KO cells cluster separately at basal but converge after LPS treatment (Fig. 6a). The top three metabolites that are increased in KO monocytes are two long chain unsaturated fatty acids-linoleic and oleic acids, and stigmasterol (Fig. 6b). This is coherent with pathway analysis by MetaboAnalyst, which showed perturbations to lipid metabolism in *C15orf48-*KO cells (Fig. 6c). Interestingly, there is a specific increase in long chain fatty acids (LCFA) in the KO cells at basal, compared to the WT cells (Fig. 6d). scRNA-seq of the monocytes show no difference in CD36 expression, indicating that this increase in LCFA is not due to elevated cellular import (Suppl. Fig. 6a). The plasma levels of LCFAs are also not elevated in *C15orf48-*KO mice (Suppl. Fig. 6b).

**Figure 6.**
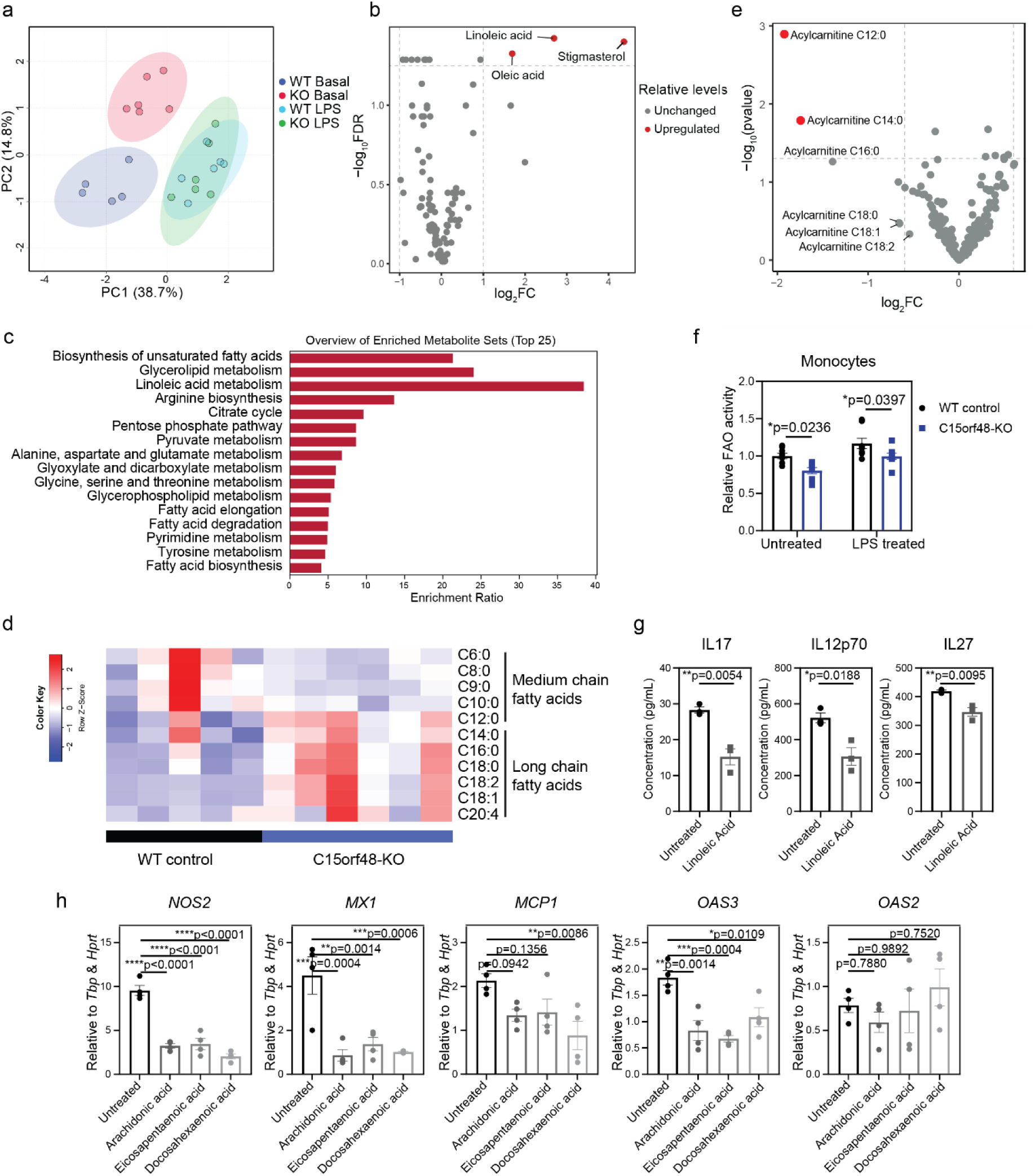
Reduced FAO in *C15orf48*-KO monocytes is associated with reduced monocyte activation. (a) PCA plot of the first two principal components of the targeted metabolomics on WT and *C15orf48-*KO monocytes treated with PBS or LPS for 4 hours. PCA plot was generated by MetaboAnalyst. n = 6 biological replicates for all, except WT basal, where n = 5. Each biological replicate contained cells pooled from 3-4 mice. (b) Volcano plot of the targeted metabolomics on WT and *C15orf48-*KO monocytes at basal. Log_2_ fold change (*C15orf48-*KO/WT) for each detected metabolite are plotted. N = 6 biological replicates for *C15orf48-*KO and 5 biological replicates for WT. Each biological replicate contained cells pooled from 3-4 mice. P-adjusted is calculated by the Benjamini-Hochberg procedure. (c) Pathway enrichment analysis generated by MetaboAnalyst on the targeted metabolomics of WT and *C15orf48-*KO monocytes at basal. (d) Relative levels of long chain and medium chain fatty acids in WT and *C15orf48-*KO monocytes at basal, based on the targeted metabolomics. n = 6 biological replicates for *C15orf48-*KO and 5 biological replicates for WT. Each biological replicate contained cells pooled from 3-4 mice. (e) Volcano plot of targeted lipidomics done on WT and *C15orf48-*KO monocytes at basal. n = 6 biological replicates. Each biological replicate contained cells pooled from 3-4 mice. P-adjusted is calculated by the Benjamini-Hochberg procedure. (f) Quantification of fatty acid oxidation activity on WT and *C15orf48-*KO monocytes at basal and after 6h LPS treatment. n = 7 biological replicates over three repeats. Each biological replicate contains cells pooled from 2-3 mice. (g) Cytokine secretion from monocytes treated with 50 µM linoleic acid. P = two-tailed unpaired Student’s t test; n = 3 biological replicates. (h) qPCR of proinflammatory genes in monocytes treated with 20 µM arachidonic acid, 50 µM eicosapentaenoic acid and 50 µM docosahexaenoic acid.

To understand the effects of the observed accumulation in LCFA and the fate of these fatty acids in the context of inflammation, we performed lipidomics on WT and *C15orf48-*KO monocytes. While there were no differences in ceramides and spermidines, which are lipids with prominent roles in macrophage activation^38^, we observed a reduction in long chain acylcarnitines (Fig. 6e, Suppl. Table 2). Carnitines are attached to fatty acids for their import into the mitochondria and subsequent beta-oxidation. Lower levels of long chain acylcarnitines suggest reduced mitochondrial import of LCFAs and reduced fatty acid oxidation (FAO), leading to the observed LCFA accumulation in KO monocytes. Indeed, using a radioactive palmitate oxidation assay, we found that FAO activity is lower in *C15orf48-*KO monocytes (Fig. 6f). These results suggest that reduced mitochondrial import and oxidation of LCFAs lead to their accumulation. While reduced FAO has been linked to increased M1 polarization and increased BMDM activation, our results here suggest that the opposite is true for monocytes in vivo^39^. We therefore sought to clarify the downstream mechanism. LCFAs such as linoleic acid and oleic acid are precursors to eicosanoids that induce anti-inflammatory response in macrophages^40,41^. We hypothesized that their accumulation might have led to increase in anti-inflammatory eicosanoid production^42,43^. We first confirmed that linoleic acid suppressed monocyte activation, as measured by their reduced production of pro-inflammatory cytokines after LPS stimulation (Fig. 6g). Lipid mediator profiling of *C15orf48*–KO monocytes revealed increased levels of arachidonic acid–derived metabolites, but not linoleic acid–derived mediators (Suppl. Fig. 6c,d, Suppl. Table 3). As linoleic acid is a metabolic precursor of arachidonic acid via sequential desaturation and elongation^44^, accumulated linoleic acid can be further converted into arachidonic acid, suggesting that the accumulated linoleic acid in KO cells is channelled into arachidonic acid. Consistent with this, treating WT monocytes with arachidonic acid, eicosapentaenoic acid, and docosahexaenoic acid suppress pro-inflammatory gene expression after LPS treatment, indicating that accumulation of these polyunsaturated fatty acids is anti-inflammatory (Fig. 6h).

Altogether, these data identify impaired FAO and the consequent buildup of anti–inflammatory polyunsaturated fatty acids and their mediators as one major mechanism by which *C15orf48* deficiency dampens monocyte activation.

### *C15orf48* supports BCAA catabolism essential for monocyte activation

In addition to changes in lipid levels, we also observed significant accumulation of some amino acids in *C15orf48-*KO monocytes (Fig. 7a), that were not due to differences in plasma amino acid levels (Suppl. Fig. 7a, Suppl. Table 4). Notably, the branched chain amino acids (BCAA) – leucine, isoleucine and valine – were all increased in the KO group. Compared to the other amino acids, BCAAs and their intermediates have bioactive properties beyond protein synthesis^45^. BCAAs are converted to branched chain keto acids (BCKAs) by BCAT1 in the cytoplasm and BCAT2 in the mitochondria (Fig. 7b). In the mitochondria, the BCKAs are then metabolized to byproducts such as propionyl CoA/propionylcarnitine and citric acid cycle products (Fig. 7b). Inhibition of BCAA catabolism has been reported to blunt macrophage and T cell response^46,47^. To determine if BCAAs are involved during monocyte activation, we added labelled α-keto-β-methylvaleric acid (KMV), the ketoacid produced from isoleucine, and quantified the levels of labelled propionylcarnitine and isoleucine with and without LPS treatment. LPS increased the amount of labelled propionylcarnitine but not isoleucine, indicating that BCAA catabolism is increased after activation (Suppl. Fig. 7b). Inhibition of BCAA uptake with 2–amino–2–norbornanecarboxylic acid (BCH) reduced cytokine secretion by monocytes, indicating that BCAA availability is required for full monocyte activation (Fig. 7c). After BCAAs are converted to BCKAs, the rate–limiting enzyme BCKDH (Branched-Chain α-Keto Acid Dehydrogenase complex) catalyzes their oxidative decarboxylation, and is inhibited by phosphorylation via BDK (Branched-chain ketoacid dehydrogenase kinase). Thus, inhibiting BDK with 3,6-dichorobenzo[b]thiopene-2-carboxylic acid (BT2)^48^ would be expected to enhance BCAA catabolism and augment activation. Unexpectedly, BT2 also reduced pro–inflammatory cytokine secretion (Fig. 7c). Because BCH limits BCAA uptake whereas BT2 drives BCKA consumption, their only shared effect is to lower intracellular BCKA levels, and both interventions blunt monocyte activation. This convergence suggested that BCKAs themselves might be the key activators, rather than BCAAs *perse*. To test this, we reduced extracellular BCAAs to a concentration that allowed normal protein synthesis (80 nM) and supplemented monocyte cultures with excess BCAAs or BCKAs. Under these conditions, only BCKA supplementation increased pro–inflammatory cytokine secretion and reduced IL–10, supporting a model in which BCKAs promote robust monocyte activation (Fig. 7d; Suppl. Fig. 7c). Although we were unable to directly quantify BCKAs in WT and *C15orf48*–KO monocytes because of their rapid turnover and limited cell numbers, genetic disruption of the mitochondrial BCAA transporter SLC25A44 markedly reduced cytokine secretion by monocytes (Fig. 7e). BCAA tolerance test showed that BCAA levels are indeed elevated in *Slc25a44*-KO mice, indicating disrupted BCAA metabolism (Suppl. Fig. 7d). As *Slc25a44*-KO monocytes still retain cytosolic BCAAs for protein synthesis, their reduced activation further implies that mitochondrial BCAA/BCKA metabolism serves a signaling role in monocyte activation beyond simple support of translation.

**Figure 7.**
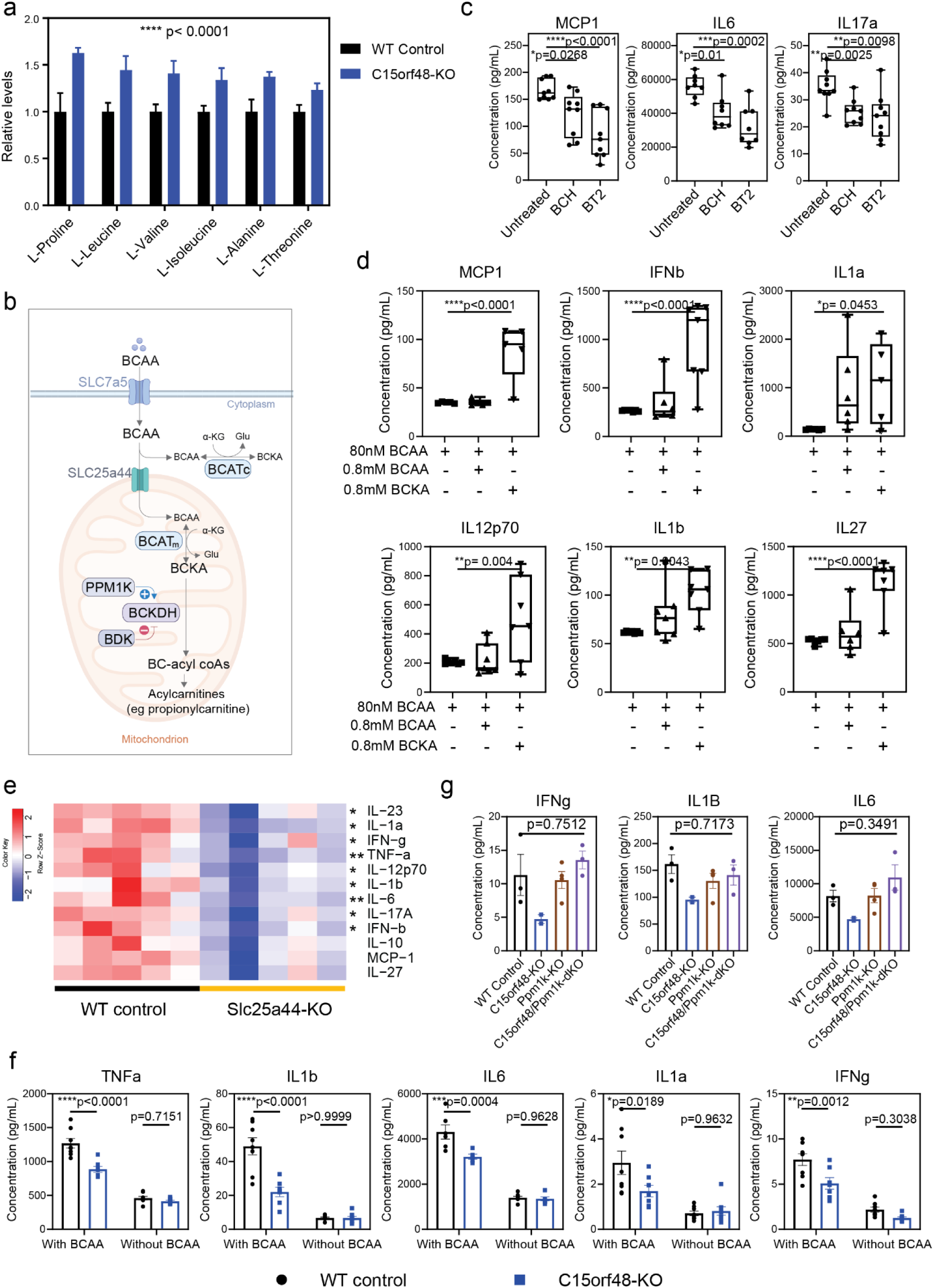
Reduced BCAA metabolism in *C15orf48*-KO monocytes is associated with reduced monocyte activation. (a) Levels of amino acids that are different between WT and *C15orf48-*KO monocytes, as measured by targeted metabolomics. P = two-way ANOVA; n = 4 biological replicates per condition from two repeats. n = 6 biological replicates for *C15orf48-*KO and 5 biological replicates for WT. Each biological replicate contained cells pooled from 3-4 mice. (b) Schematic of BCAA metabolism in cells. Created in https://BioRender.com. (c) Cytokine secretion after LPS from WT monocytes treated with 10mM leucine transporter inhibitor (BCH) and 100 µM BDK inhibitor (BT2). (d) Cytokine secretion after LPS from WT monocytes cultured in 80nM BCAA with 80mM BCAA or 80mM BCKA. P = one-way ANOVA; n = 6 biological replicates per condition from two repeats. (e) Quantification of cytokines secreted by WT and *Slc25a44*-KO monocytes treated with LPS for 6 hours. n = 10 biological replicates over two repeats. Only 1 repeat is shown here. P = two-tailed unpaired Student’s t test based on all 10 biological replicates. (f) Response of WT and *C15orf48-*KO monocytes to LPS, with and without BCAA in the media. P = two-way ANOVA; n = 6 biological replicates per condition from two repeats. (g) Cytokine secretion from WT, *C15orf48-*KO, *Ppm1k*-KO, *Ppm1k*/*C15orf48-*dKO monocytes after LPS treatment. P = one-way ANOVA; n = 3 biological replicates for WT, *Ppm1k*-KO, *Ppm1k*/*C15orf48-*dKO; 2 biological replicates for *C15orf48-*KO.

Having established that BCAAs are involved in monocyte activation, we checked whether altered BCAA metabolism in *C15orf48*–KO monocytes could account for the suppressed activation. WT and *C15orf48-*KO monocytes were activated with LPS in the presence and absence of BCAAs. As expected, in BCAA–replete conditions, WT monocytes secrete more cytokines than *C15orf48-*KO monocytes, whereas removing BCAAs largely eliminated this, indicating that BCAAs are necessary to reveal the activation defect of *C15orf48*–KO monocytes (Fig. 7f).

Our results suggest that increasing BCKA levels in *C15orf48-*KO monocytes might restore their response to LPS. Protein phosphatase, Mg^2+^/Mn^2+^ dependent 1K (*Ppm1k*) is a phosphatase that activates BCKDH. Knocking out *Ppm1k* decreases BCKDH activity, which decreases BCAA catabolism^49^ and is expected to cause BCKA accumulation. Consistent with this prediction, *Ppm1k*–deficient mice showed higher *Cox2* and *Cxcl1* transcript levels in the lungs after LPS injection (Suppl. Fig. 7e)., although plasma cytokines were not increased, likely due to systemic compensatory mechanisms that maintain overall BCAA metabolism (Suppl. Fig. 7f)^50^. To bypass such whole-body compensation, we isolated monocytes from WT, *C15orf48-*KO, *Ppm1k*-KO and *C15orf48/Ppm1k*-double KO (dKO) mice and treated them with LPS. Whereas *C15orf48-*KO cells secreted less cytokines than WT monocytes, cytokine production was restored to near–WT levels in the dKO monocytes (Fig. 7g). This data support the idea that increasing BCKA levels in *C15orf48*–deficient cells rescues their activation defect and that *C15orf48* regulates monocyte activation through sustaining BCAA/BCKA metabolism.

## Discussion

In this study, we demonstrate that *C15orf48* is upregulated in response to systemic inflammation and plays a key role in modulating host responses during Kp infection and endotoxemia. *C15orf48-*KO mice were able to suppress both Kp bacterial burden and systemic inflammation. The suppression of systemic inflammation is independent of the reduced Kp load, as LPS injection results in lower inflammation in KO mice as well. This ameliorates the survival of the KO mice in a lethal LPS challenge. In the lungs, *C15orf48* expression is induced primarily in myeloid and ECs following inflammatory stimulation. Cell-type-specific knockout experiments revealed the systemic pro-inflammatory effect of *C15orf48* is primarily driven by myeloid cells, and more specifically in monocytes. Within the myeloid compartment, no significant changes were detected in neutrophils, even though MOCCI expression is notably high in these cells. This suggests that MOCCI may play a role in neutrophils under specific conditions that we have yet to identify. *C15orf48-*KO monocytes have higher ROS and lower NAD pool, resulting in dysregulated FAO and BCAA catabolism that blunts activation.

Our observation that *C15orf48* plays contrasting roles in myeloid and ECs is supported by supported by prior studies reporting anti–inflammatory roles for *C15orf48* in the colon and ECs,^19,20^ and pro–inflammatory roles in human monocyte–derived macrophages^18,33^ and tumour–associated macrophages^21^. This supports the idea that *C15orf48* can play divergent roles in leukocytes versus non–leukocytes and more importantly, that regulation of ETC activity can yield completely different outcomes in endothelial and myeloid lineages. Our findings also differ from those of Liu et al. (2023), who reported reduced survival and higher pathogen burden in *C15orf48*–KO mice infected with influenza, Kp *or Staphylococcus aureus*^51^. They proposed that MOCCI binds and inhibits gasdermin D (GSDMD) activation to limit pyroptosis, although this mechanism is difficult to reconcile with MOCCI’s mitochondrial localization. While we cannot fully resolve these discrepancies, differences in infection models and pathogen strains, including the Kp strain used, are likely to contribute. Differing infection models or strains that have greater vascular activation might tip the balance towards anti-inflammatory function of *C15orf48* in ECs when using whole-body KOs. Altogether, these studies highlight that the *C15orf48*–CIV–miR–147b axis can have distinct, even opposing, consequences depending on tissue and context, making it essential to dissect its function in each relevant cell type rather than inferring a uniform role across tissues.

Our study demonstrates that *C15orf48* is required by monocytes to maintain a bioenergetic and metabolomic state that is poised for optimal activation following an inflammatory challenge. *C15orf48-*KO monocytes have decreased mitochondrial respiration and FAO, and accumulation of LCFAs and BCAAs. We propose that surplus LCFAs are in turn channelled towards the production of anti-inflammatory lipid mediators, while reduced BCAA catabolism limits the availability of pro-inflammatory BCKAs, and these two pathways together limit pro-inflammatory cytokine secretion in KO monocytes. How *C15orf48* deficiency, ETC impairment, reduced BCAA catabolism and reduced FAO are connected is still mechanistically unclear. We propose the following scenario. *C15orf48* deficiency affects CIV activity and ETC efficiency. This can directly and indirectly reduce NAD^+^ redox potential and overall availability. Beyond NAD’s role as an electron carrier in several metabolic pathways where it cycles between an oxidised and reduced state, NAD is also consumed in several cellular processes, such as PARP–mediated DNA repair, sirtuin–dependent deacylation, and CD38–driven NADase activity^52^. It is replaced through NAMPT-dependent salvage pathway that recycles nicotinamide and *de novo* synthesis from tryptophan. Macrophages deplete NAD^+^ upon activation, due to high CD38, PARP and sirtuin 3 activity^36,53^. Despite increasing the activity of the salvage pathway to compensate for the loss of NAD^+^, the total NAD^+^ pool still shrinks after the macrophage encounter an inflammatory trigger^36^. *C15orf48-*KO monocytes, owing to ETC impairment and potentially higher DNA damage induced by increased mitochondrial ROS, have a smaller NAD^+^ pool to buffer against the LPS-induced NAD^+^ consumption. Disruption of NAD^+^ homeostasis, in turn, impairs FAO and BCAA catabolism, both of which rely on NAD–dependent dehydrogenase reactions (β–hydroxyacyl–CoA dehydrogenase and BCKDH respectively), leading to LCFA and BCAA accumulation. We therefore propose that *C15orf48* regulates ETC efficiency and mitochondrial ROS production, which in turn preserves NAD^+^ levels to allow for robust monocyte activation. Upon activation, monocytes are required to adapt rapidly to changing metabolic demands, switching between pathways such as glycolysis, BCAA and fatty acid metabolism. By supporting NAD^+^ homeostasis and related metabolic processes, *C15orf48* facilitates the ability of monocytes to efficiently reprogram their metabolism in response to immune challenges. How MOCCI induction in monocytes remodels CIV to maintain ETC efficiency and suppress mitochondrial ROS following an inflammatory trigger - in the absence of a measurable effect on CIV activity - is a subject of future research.

Our BALF inoculation assay reveals that the lung environment provides nutrients conducive to Kp growth, with further enhancement observed in BALF from infected WT mice. Notably, while BALF from uninfected WT and *C15orf48-*KO mice shows no difference, BALF from infected *C15orf48-*KO mice fails to support increased Kp growth to the levels of infected WT mice. The more inflammatory milieu in the WT lungs could potentially be more conducive for Kp growth, as Kp utilizes host-derived proinflammatory metabolites and cytokines to aid in their colonization^30,31^. Certain virulent clinical isolates of Escherichia coli can even bind human cytokines and use them as growth–promoting factors^54^. BALF from influenza A virus–infected lungs has also been reported to enhance the growth of secondary bacterial pathogens in vitro^55^. Besides the direct effect of pro-inflammatory mediators on bacterial growth, an alternative possible explanation is that the reduced inflammatory response in *C15orf48-*KO mice lowers vascular permeability compared to WT mice, thereby limiting nutrient leakage into the alveoli. Another hypothesis is that Kp thrives on metabolites such as succinate and itaconate, which are produced by activated cells^56,57^. The lower inflammatory state in *C15orf48-*KO mice leads to reduced production of these metabolites, thus limiting their availability to Kp. Additionally, we cannot dismiss the possibility that *C15orf48-*KO mice may secrete an inhibitory factor that restricts Kp proliferation, whose identity is beyond the scope of this study.

The observed beneficial effects of *C15orf48-*KO on bacterial load and survival appear counter-intuitive at first glance. This raises the question why would *C15orf48* evolve if its absence seems advantageous? As our models mimic late stages of infection where there are high bacterial load and systemic inflammation, the KO of *C15orf48* proves beneficial in these models as it mitigates excessive inflammation. However, at early stages of infection, strong myeloid cell activation is required for rapid pathogen clearance. *C15orf48* would therefore be crucial in robust monocyte activation to clear infections before they spread.

Altogether, our study presents *C15orf48* acts as a key metabolic checkpoint in myeloid cells, enabling robust monocyte activation and proinflammatory responses through regulation of mitochondrial respiration, fatty acid oxidation, and BCAA metabolism. The observation that *C15orf48-*KO mice exhibit enhanced resistance to infection and sepsis despite reduced inflammation challenges the traditional view that a heightened immune response is always beneficial for pathogen clearance, and instead points towards a model where metabolic fine-tuning of immune cells can promote effective host defense while limiting immunopathology. Targeting the *C15orf48* or the downstream BCAA/FAO metabolic pathways may offer novel therapeutic strategies to modulate monocyte-driven inflammation in a range of infectious and inflammatory diseases, potentially mitigating tissue damage and improving patient outcomes.

## Acknowledgements

This work was supported by the National Medical Research Council, Singapore (MOH-000632 awarded to C.L.; MOH-000960 and MOE-T2EP30122-0011 awarded to L.H.). Y.W. was supported by Tanoto Foundation in Diabetes. This research used NCRIS-enabled Metabolomics Australia infrastructure at the University of Melbourne and funded through BioPlatforms Australia. We gratefully acknowledge the Duke-NUS Metabolomics Facility and NUS SLING for their services. BCAA flux analyses were performed by the Duke Molecular Physiology Institute metabolomics core with funding from NIH grant P30DK124723. We also extend our appreciation to Chao Liang, Shuang Wang, and Jiemin Nah for their technical assistance and providing essential reagents.

## Declaration of generative AI and AI-assisted technologies in the writing process

During the preparation of this work, the authors used ChatGPT in order to improve the readability of some sentences. After using this tool/service, the authors reviewed and edited the content as needed and take full responsibility for the content of the published article.

## Methods

### Mouse generation and maintenance

All animal protocols (2017/SHS/1336 and 2017/SHS/1767) were approved by the NUS Institutional Animal Care and Use Committee (IACUC). *C15orf48*-fl/fl mice were generated by the Melbourne Advanced Genome Editing Centre in the Walter and Eliza Hall Institute of Medical Research. LoxP sites were inserted upstream of exon 2 and downstream of exon 4 of *AA467197*, the mouse ortholog of *C15orf48*. The mice were mated with ROSA26-creERT2^1^, VE-cad-creERT2 (Taconic Biosciences 13073) and Lysm-cre mice^2^ to generate cell-type specific knockout mice. Mice carrying cre or creERT2 transgenes were maintained as heterozygotes, while floxed alleles are maintained in the homozygous state. Mice with creERT2 were oral gavaged with 50 g tamoxifen per kg of mouse weight (Merck T5648) for 4 consecutive days. They were used for experiments after 1 week of resting period for the tamoxifen to flush out from them. *Slc25a44*-KO were obtained by crossing *Slc25a44*-fl/fl mice with Zp3-Cre transgenic mice to generate germline deletion of *Slc25a44*. *Slc25a44*-fl/fl mice were generated at the Transgenic and Reproductive Technology Centre of the National University of Singapore. LoxP sites were inserted up and downstream of exon 2 of *Slc25a44*. *Ppm1k*-KO mice were generated and maintained as previously reported^3^. All experimental mice were maintained on a C57BL/6J background.

### Quantification of mouse metabolism

Mouse temperature was measured using a rodent probe (BIOSEB Lab BIO-TK885) inserted into the anus of the mouse. Mouse activity, oxygen consumption, carbon dioxide production, food and water intake were concurrently monitored using the PhenoMaster automated home-cage system (TSE Systems, Germany). Briefly, mice were singly housed in metabolic cages equipped with infrared beam arrays to quantify locomotor activity. The system continuously measured oxygen and carbon dioxide concentrations in the cage, allowing calculation of energy expenditure and respiratory exchange ratio (RER) as previously described. Food and water intake were automatically recorded via high-precision balances and drinking sensors integrated into each cage. Animals were acclimatized to the system for 24 h prior to data collection. Measurements were collected over 24 h before LPS injection and for 16h after LPS injection. Environmental conditions were maintained at 24°C, 12h light/dark cycles.

### Infection of mice with *Klebsiella pneumoniae*

*Klebsiella pneumoniae* strain MGH78578 (ATCC 700721) from -80°C frozen stock was thawed and inoculated in Luria-Bertani (LB) Broth (Miller) (Axil Scientific BIO-4000), then cultured at 37°C with shaking at 160 rpm overnight. Bacterial cultures were diluted 1:50 into fresh LB Broth the following day and grown for 4 h. Bacteria were pelleted at 6000 g for 10 minutes at room temperature and washed with sterile phosphate-buffered saline (PBS) twice. Concentration of inoculum was quantified using optical density measurement method at 600 nm (OD 600nm) and further confirmed by plating onto LB agar plate (Axil Scientific BIO-4010) after serial dilution to quantify colony forming units (CFUs).

Mice were anesthetised with for ketamine (20mg/kg body weight) and xylazine (2mg/kg body weight) cocktail by intraperitoneal injection and infected intranasally with 8x 10^8^ CFU of *K. pneumoniae* in 40 µL of sterile PBS. Control was treated with 40 µL of sterile PBS alone. The mice were monitored daily for clinical signs of distress and changes in body weight after infection. Body temperatures of mice were taken 1 week prior and at indicated time after infection. The mice were sacrificed at 18 h and 42 h post infection.

### Processing of mouse tissue for assays

Bronchoalveolar lavage fluid (BALF) was collected from the lungs by flushing the lungs with 1 mL of PBS through the trachea. For cytokine analysis, in vitro *K. pneumoniae* growth assays and metabolomics analysis, the BALF was centrifuged at 500 g for 5 minutes at 4°C and the supernatant was collected and stored at -80°C. Single cells from lungs were obtained by mincing the lungs and incubating them in dissociation medium for 30 minutes at 37°C, followed by homogenisation using gentleMACS^TM^ Dissociators (Miltenyi Biotec 130-134-029). This was followed by washing in 2% FCS/PBS, supplemented with 1mM EDTA. The cells were strained through 100 µm cell strainers (SPL Life Sciences 93100), followed by 70 µm cell strainers (SPL Life Sciences 93070).

Blood was collected into tubes coated with EDTA and spun down at 2500 g for 10 minutes at 4°C. The plasma was aliquoted and at -80°C for further analysis.

### Quantification of *K. pneumoniae* burden

Liver and spleen were weighed and homogenised using the plunger of a 3 mL syringe through a 70 µm cell strainer (SPL Life Sciences 93070) in sterile PBS with 0.2% Tween® 20 (Sigma P1379). An aliquot of BALF, blood, liver and spleen homogenates were serially diluted in sterile PBS and plated on LB agar plates in duplicates for the enumeration of *K. pneumoniae* colonies.

### In vitro *K. pneumoniae* growth assay in BALF

Aliquots of frozen BALF supernatants from uninfected mice and *K. pneumoniae* infected mice were thawed on ice and inoculated with K. Pneumoniae at a final concentration of 9300 CFU/mL. The mixtures were incubated at 37°C with shaking at 1100 rpm for 2 h and 4 h and then serially diluted and plated on LB agar plates for quantification. Un-inoculated BALF supernatant from infected mice were also plated on LB plates to control for residual *K. pneumoniae*.

### Monocyte, macrophage and neutrophil isolation from mouse bone marrow

Mice were euthanized by CO_2_ inhalation and femurs and tibias were surgically isolated. The bones were disinfected in 70% ethanol for 5 mins and rinsed twice with sterile phosphate-buffered saline (PBS, Cytiva SH30028.02). Bone marrow was flushed out with cold RPMI-1640 (Cytiva SH30027.01) with 25G needle attached to a syringe and strained through 70 μm cell strainer (SPL Life Sciences 93070). The cell suspension was pelleted, resuspended and incubated in red blood cell lysis buffer (BioLegend 420301) for 3 minutes, followed by neutralisation with PBS containing 2% heat-inactivated fetal bovine serum (FBS v/v, Cytiva SV30160.03HI) and 1mM EDTA. The cells were pelleted again and resuspended in desired medium.

Murine monocytes and neutrophils were isolated from mouse bone marrow using EasySep™ Mouse Monocyte Isolation Kit ( STEMCELL Technologies 19861) and MojoSort™ Mouse Ly-6G Selection Kit (BioLegend 480124). LS Columns (Miltenyi Biotec 130-042-401) were used with the kits, according to the manufacturer’s protocol. Monocytes were cultured in 10% FCS/DMEM supplemented with 25 ng/mL M-CSF (Thermo Fisher Scientific PMC2044).

To derive mouse BMDMs, the isolated bone marrow was seeded into L929-conditioned media for 7 days. The media was changed every 3 days. After 7 days, the cells were scraped and reseeded at 50,000 cells/cm^2^ for experiments.

### Isolation of human monocytes and monocyte-derived macrophages

PBMCs were isolated from blood of healthy donors using density gradient centrifugation (Sigma Histopaque®-1077). Human CD14^+^ monocytes were separated negatively from peripheral blood mononuclear cells (PBMCs) by EasySep™ Direct Human Monocyte Isolation Kit (STEMCELL Technologies 19669) and LS Columns (Miltenyi Biotec 130-042-401). To derive MDMs, PBMCs from 7.5 mL of blood cones were seeded into T175 flask in 10% human serum/RPMI, supplemented with 100 ng/mL macrophage colony-stimulating factor (M-CSF) for 7 days at 37°C with 5% CO₂. The medium was replaced every 3 days. After 7 days of differentiation, the resulting MDMs were harvested by scraping and re-seeded for experiments.

### Quantification of mitochondrial membrane potential and ROS levels in monocytes

Monocytes were isolated from the bone marrow and cultured overnight in 10% FCS/DMEM supplemented with 25 ng/mL M-CSF (Thermo Fisher Scientific PMC2044) in 1.5 mL Eppendorf tubes with open lids. The next morning, the cells were pelleted at 500 g for 5 min at rtp. They were then resuspended in warm media containing the dye and incubated at 37°C for 30 min. The concentrations of the dyes used were 100 nM and 5 µM for TMRE and mito-SOX, respectively. The dye intensity of the cells was then acquired on a flow cytometer.

### Collection of monocytes for metabolomics analysis

Monocytes were isolated from bone marrow. 2.5 million monocytes per condition were cultured in 100 μL of media in 1.5 mL Eppendorf tubes. The media consist of 10% FCS/DMEM/25 ng/mL M-CSF, with 100 ng/mL LPS added in the indicated conditions. The cells were cultured at 37°C for 4 hours. 400 μL of ice cold PBS was added to the cells to stop all enzymatic activities. The cells were pelleted at 500 g for 5 min at 4°C. The supernatant was collected for cytokine assays while the cell pellet was washed two more times with ice cold PBS. The cells were then snap frozen in liquid nitrogen. For the BCAA flux analysis, 1mM of ¹³C₆ α-keto-β-methylvalerate (Cambridge Isotope Laboratories CLM-8669) was spiked into the media and the incubation was lengthened to 6 hours to allow accumulation of the downstream labelled metabolites.

### BCAA tolerance test

BCAA solutions were freshly prepared in sterile 0.9% saline and contained 82.5 mM leucine, 37.5 mM isoleucine, and 60 mM valine, corresponding to a molar ratio of 2:1:1.5 (leucine:isoleucine:valine). Male WT control and *Slc25a44*-KO mice (10–14 weeks of age) were fasted for 6 h (9am–3pm) with free access to water. Following the fasting period, mice received an intraperitoneal injection of the BCAA solution at a dose of 8 μL/g body weight. Blood samples were collected via retro-orbital bleeding immediately before injection (0 min) and at 30 min after BCAA administration. Blood samples were allowed to clot on ice for 30 min and centrifuged at 10,000 g for 10 min at 4°C. Serum was collected and stored at −80°C until analysis by liquid chromatography–mass spectrometry (LC–MS).

### Gas chromatography mass spectrometry for polar metabolites from mouse monocytes

Polar metabolites were extracted by adding 200 µL of extraction solvent (3 parts methanol:1 part water). The extraction solvent was prepared by premixing 3 mL methanol, 980 µL water, 10 µL of 1 mM ^13^C₆-sorbitol, and 10 µL of 1 mM ^13^C₅,^15^N-valine. Samples were vortexed vigorously and incubated on ice for 15 min. The samples were vortexed every 5 min to enhance extraction. After a second vigorous vortex at the end of the 15 min incubation, samples were centrifuged at 16,000 g for 5 min at 4 °C. The resulting supernatant was carefully transferred to a new 1.5 mL microcentrifuge tube and snap-frozen in liquid nitrogen for subsequent processing by Metabolomics Australia.

To prepare a pooled biological quality control (PBQC) sample, 75 µL from each extract was combined. 100 µL of each study sample and the PBQC were transferred into HPLC inserts and evaporated at 30 °C to complete dryness, using a CHRIST RVC 2-33 CD plus speed vacuum. To limit the amount of moisture present in the insert, 50 µL 100% methanol (LCMS grade) was added to each insert and evaporated using a speed vacuum.

Dried samples for GCMS analysis were derivatised online using the Shimadzu AOC6000 autosampler robot. Derivatisation was achieved by the addition of 25 µL methoxyamine hydrochloride (30 mg/mL in pyridine, Merck) followed by shaking at 37 °C for 2 h. Samples were then derivatised with 25 µL of N,O-bis (trimethylsilyl)trifluoroacetamide with trimethylchlorosilane (BSTFA with 1% TMCS, Thermo Scientific) for 1 h at 37 °C. The sample was allowed to equilibrate at room temperature for 1 h before 1 µL was injected onto the GC column using a hot needle technique. Split (1:10) injections were performed for each sample.

The GC-MS system used comprised of an AOC6000 autosampler, a 2030 Shimadzu gas chromatograph and a TQ8050NX triple quadrupole mass spectrometer (Shimadzu, Japan) with an electron ionisation source(−70eV). The mass spectrometer was tuned according to the manufacturer’s recommendations using tris-(perfluorobutyl)-amine (CF43). GC-MS was performed on a 30m Agilent DB-5 column with 0.25 mm internal diameter column and 1 µm film thickness. The injection temperature (inlet) was set at 280 °C, the MS transfer line at 280 °C and the ion source adjusted to 200 °C. Helium was used as the carrier gas at a flow rate of 1 mL/min and argon gas was used in the collision cell to generate the MRM product ion. The analysis of the derivatised samples was performed under the following oven temperature program; 100 °C start temperature, hold for 4 minutes, followed by a 10 °C/min oven temperature ramp to 320 °C with a following final hold for 11 minutes.

Samples were analysed in a randomised order, with a pooled biological quality control sample added after every 5 samples. The pooled biological quality control was used to monitor downstream sample stability and analytical reproducibility. Approximately 500 targets were collected using the Shimadzu Smart Metabolite Database, where each target comprised a quantifier MRM along with a qualifier MRM, which covers approximately 356 unique endogenous metabolites and multiple stable isotopically labelled internal standards. Resultant data was processed using Shimadzu LabSolutions Insight software, where peak integrations were visually validated and manually corrected where required.

### Targeted lipidomics on mouse monocytes

Targeted lipidomics on ceramides, spermidines and acylcarnitines were carried out on 2.5 million monocytes as previously described^4^. Lipids were extracted from cell pellets using a single-phase butanol:methanol (1:1, v/v) including internal standards. The standards were purchased from Avanti Lipids and included acylcarnitine 16:0 D3, cholesterol ester 18:0 D6, dihydroceramide d18:0/08:0, ceramide d18:1/12:0, deoxyceramide m18:1/12:0, diacylglycerol 15:0/15:0, GM3 d18:1/18:0 D3, monohexosylceramide d18:1/12:0, dihexosylceramide d18:1/12:0, trihexosylceramide d18:1/18:0 D3, lysophosphatidylcholine 13:0, lysophosphatidylethanolamine 14:0, phosphatidylcholine 13:0/13:0, plasmalogen phosphatidylcholine 18:0/18:1 D9, phosphatidylethanolamine 17:0/17:0, plasmalogen phosphatidylethanolamine 18:0/18:1 D9, phosphatidylglycerol 17:0/17:0, phosphatidylinositol 12:0/13:0, phosphatidylserine 17:0/17:0, sphingomyelin d18:1/12:0, sphingosine d18:1 D7, triacylglycerol 12:0/12:0/12:0. Phosphatidylglycerol 17:0/17:0 was used for normalisation of the signal of BMP in the samples. The mixture was vortexed for 10 seconds, sonicated for 30 min and then centrifuged at 4°C (14,000 g for 10 min). The supernatant fraction was collected for LC-MS/MS analysis. A pooled lipid extract was used as a quality control (QC) sample and injected every five study samples. Data were normalized to protein concentration in each sample. The LC-MS/MS analysis was performed on an Agilent UHPLC 1290 Infinity II liquid chromatography system connected to an Agilent QqQ 6495C. An Agilent Zorbax RRHD Eclipse Plus C18 column (2.1 × 50 mm, 1.8 μm) was used for the RPLC separation. The mobile phases A (60% water and 40% acetonitrile with 10 mmol/L ammonium formate) and B (10% acetonitrile and 90% isopropanol with 10 mmol/L ammonium formate) were used for the chromatographic separation. The following gradient was applied: 0-2 min, 20-60% B; 2-12 min, 60-100% B; 12-14 min, 100% B; 14.01-15.8 min, 20% B. The oven temperature was maintained at 40°C. Flow rate was set at 0.4 mL/min and the sample injection volume was 2 or 5 μL, depending on the sample type. The positive ionization spray voltage and nozzle voltage were set at 3,000 V and 1,000 V, respectively. The drying gas and sheath gas temperatures were both maintained at 250 oC. The drying gas and sheath gas flow rates were 14 L/min and 11 L/min, respectively. The nebulizer nitrogen gas flow rate was set at 35 psi. The iFunnel high and low pressure RF were 150 V and 60 V, respectivelyData were analysed with Agilent MassHunter Quantitative Analysis (v8). Lipids were identified by retention time and MRM transitions, with quantitation based on normalization of peak areas to class-specific internal standards and total protein content. Only lipids reproducibly detected (CV < 30%) and not found in blank controls (<10% of QC) were quantified. Relative quantitation was performed using a single standard per lipid class.

### Quantification of amino acids in mouse plasma

A volume of 50 µl from each sample was combined with 15 µl of a deuterium-labeled amino acid standard mix and diluted using 400 µl of methanol. The resulting mixture was subjected to centrifugation at 13,000 rpm for 5 minutes at 4°C. The supernatant was then collected and derivatized using 3M hydrochloric acid in butanol. Liquid chromatography-mass spectrometry (LC-MS) analysis was performed with a quadrupole ion trap mass spectrometer (QTRAP 5500, AB Sciex, Olympia, DC, USA) coupled to an Agilent 1290 Infinity liquid chromatography system (Agilent Technologies, Santa Clara, CA, USA). Sample separation was achieved on a Phenomenex Luna® Omega C18 column (100 × 2.1 mm, 1.6 µm, Torrance, CA, USA). The chromatographic separation employed water (mobile phase A) and acetonitrile (mobile phase B), both containing 0.1% formic acid. The LC gradient started at 2% B for 0.8 minutes at a flow rate of 0.4 mL/min, followed by a stepwise increase to 15% B over 0.1 minutes, 20% B over the next 5.7 minutes, up to 50% B in 0.5 minutes, and finally to 70% B over another 0.5 minutes. The column was then re-equilibrated to 2% B for 0.9 minutes. Chromatographic data were processed and integrated using MultiQuant™ software version 3.0.3 (AB Sciex, DC, USA).

### Quantification of acylcarnitine in mouse plasma

For the analysis of acylcarnitines, each sample was supplemented with a deuterium-labeled acylcarnitine standard mixture and diluted with 400 µl of methanol. The mixture was centrifuged at 13,000 rpm for 5 minutes at 4°C, after which the supernatant was collected and derivatized using 3M hydrochloric acid in methanol (Sigma Aldrich). Electrospray ionization in positive mode was employed for compound ionization. The derivatized samples were analyzed using an Agilent 1290 Infinity liquid chromatography system (Agilent Technologies, CA, USA) coupled to a QTRAP 5500 quadrupole ion trap mass spectrometer (AB Sciex, DC, USA). Chromatographic separation was carried out with 80% methanol as the mobile phase at a flow rate of 0.4 mL/min. Data integration was performed with MultiQuant™ software version 3.0.3 (AB Sciex, DC, USA).

### Tracing of BCAA fluxL

2.5 million freshly isolated mouse monocytes from bone marrow were cultured in DMEM (ThermoFisher Scientific A14430) containing 2% dialyzed FBS (ThermoFisher Scientific 26-400-044), 25 ng/mL M-CSF (ThermoFisher Scientific PMC2044), 2 mM glucose, 0.4 mM L-glutamine and 1mM ^13^C_6_ sodium 2-keto-3-methyl-pentanoate (Cambridge Isotope Laboratories CILZ/CLM-8669). They were cultured with and without 100 ng/mL LPS for 6 h at 37°C. They were then washed 3 times with 1 mL of ice cold PBS and snap frozen in liquid nitrogen before they were kept at -80°C until further analysis. The quantification of labelled intermediates were carried out as previously described^5^.

### Analysis of metabolomics data

The dataset was normalised by median and log transformed. PCA and pathway analysis were generated using MetaboAnalyst 4.0^6^. R software was used to generate volcano plots^7^.

### Quantification of NAD/NADH

NAD and NADH were quantified using Q-NAD Tissue/Cell NAD+ and NADH assay kit (NADMED RUO_003), according to manufacturer’s protocol. Briefly, 2 million cells were resuspended in 300 µL of pre–warmed Extraction Buffer B (55°C) and incubated at 55°C for 2 min, and then cooled on ice. Homogenates were centrifuged at 20,000 g for 10 min at 4°C, and supernatants were collected into clean microtubes. The pellets were retained for protein quantification. The supernatants were equilibrated to room temperature. 100 µL supernatant was combined with 100 µL NAD⁺ Stabilizer and incubated for 5 min at room temperature, Another 100 µL supernatant was combined with 100 µL NADH Stabilizer, and incubated for 2 min at 75–80°C, followed by cooling on ice for 5 min. These mixtures were mixed with the Color Reagent, Assay Buffer C, and NAD Enzyme provided in the kit, incubated for 6 min in the NAD⁺ reaction and 15 min in the NADH reaction at room temperature. 10 µL Stop Solution was added to each well and absorbance was read at 573 nm.

For normalization, the protein pellets obtained after NAD⁺/NADH extraction were solubilized in 100 µL Solubilization Buffer D, followed by brief sonication. They were then quantified using BCA.

### Quantification of fatty acid oxidation (FAO)

Cell lysates prepared with 0.0025% digitonin were utilized for the fatty acid oxidation (FAO) assay. Samples were incubated in an oxidation reaction buffer containing 100 mM sucrose, 10 mM Tris-HCl (pH 8), 5 mM KH₂PO₄, 0.2 mM EDTA, 80 mM KCl, 1 mM MgCl₂, 2 mM L-carnitine, 0.1 mM malate, 0.05 mM coenzyme A, 2 mM ATP, and 1 mM DTT, in the presence of 0.7% BSA conjugated with 17 nM palmitate. For experimental conditions, 0.5 μCi of [^3^H]-palmitic acid was added. Incubations were performed at 37°C for 4 hours. Released [^3^H]H₂O was captured by adding perchloric acid and trapping the evolved gas with NaOH-soaked filter paper discs placed in Eppendorf tubes for 1 hour. The filter papers were subsequently collected for scintillation counting. Remaining reaction mixtures were centrifuged at 14,000 × g for 10 minutes, and supernatants were also analyzed by scintillation counting. Radioactivity was measured using a Tricarb scintillation counter (Perkin Elmer).

### Cytokine Analysis

Cytokine concentrations in cell culture supernatant, mouse serum, and BALF supernatant were quantified by using LEGENDplex^TM^ bead-based multiplex assay (BioLegend 740809; 740446) according to the manufacturer’s protocols. Fluorescence intensity of the beads were acquired by LSRFortessa^TM^ (BD Biosciences). The levels of cytokines were analysed using the LEGENDplex^TM^ v8.0 software.

### RT-qPCR of mRNA and miRNA

Total RNA was extracted using Quick-RNA™ MicroPrep Kit from Zymo Research (ZYR.R1051) following the manufacturer’s instructions. Cells were lysed using the lysis buffer from the kit while mouse tissues were homogenized in TRIzol™ Reagent (Thermo Fisher Scientific 15596026). For mRNA quantification, complementary DNA (cDNA) was synthesized from total RNA using iScript™ Reverse Transcription Supermix (Bio-Rad 1708841). Quantitative real-time PCR (qPCR) was performed using SYBR Green chemistry (Bio-Rad 1725125). The primers used are listed in the table below.

Mouse genes

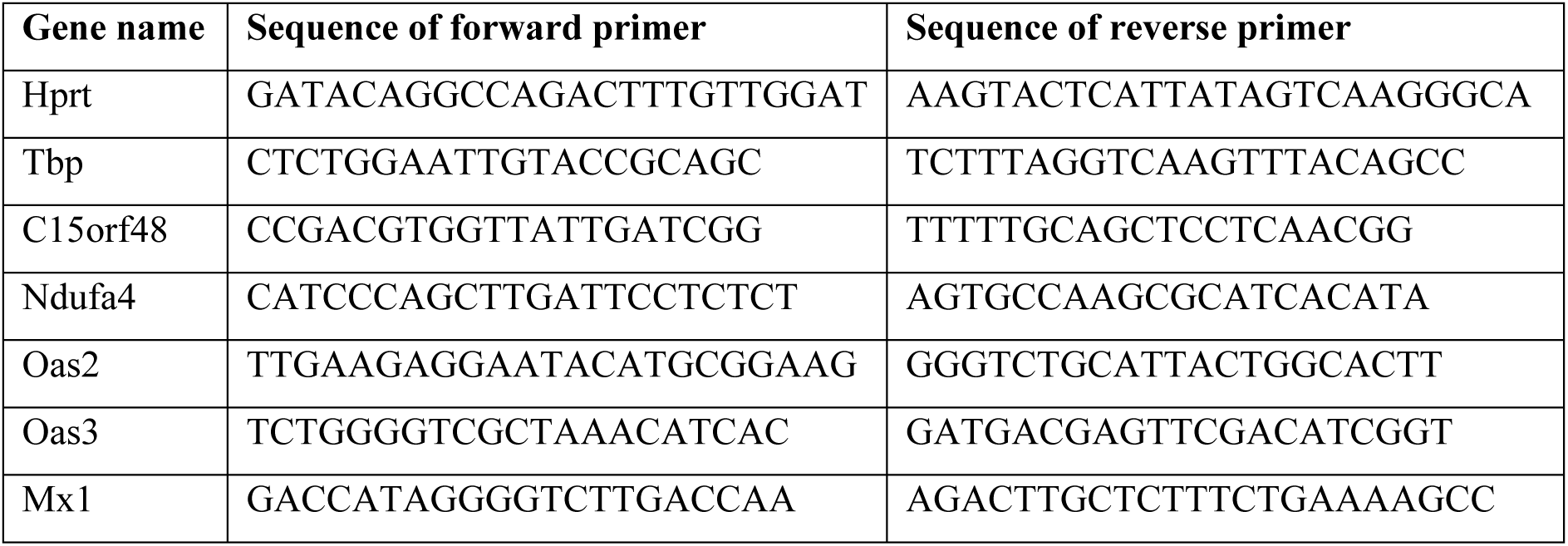
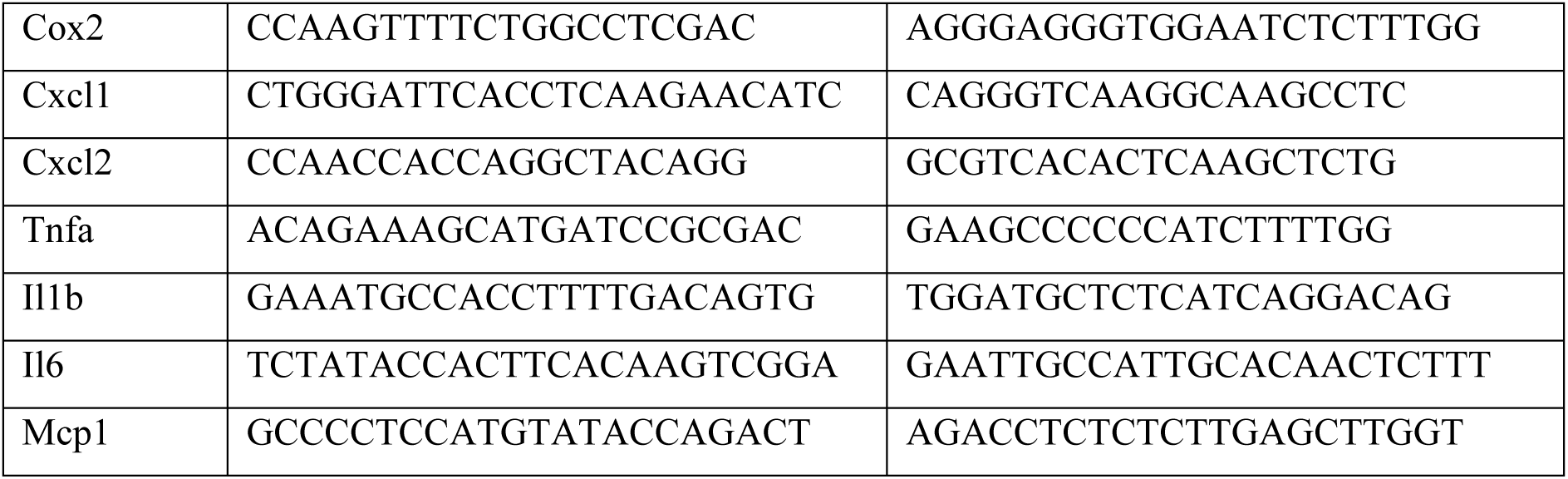

Human genes

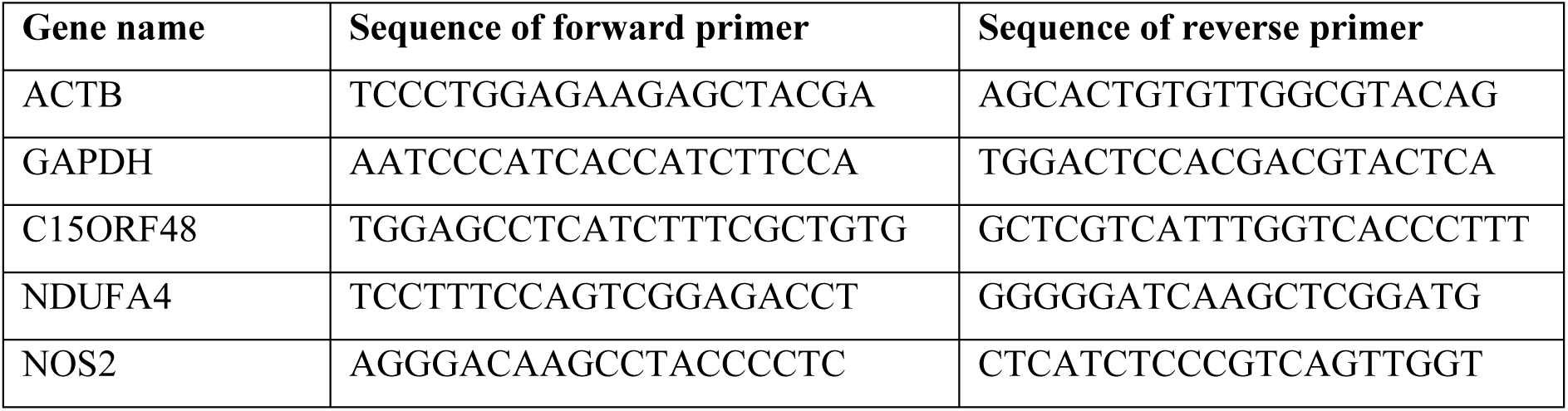

For miRNA quantification, cDNA was synthesized using the TaqMan MicroRNA Reverse Transcription Kit (Thermo Fisher Scientific 4366596) according to the manufacturer’s protocol with miRNA-specific stem-loop primers. Quantitative PCR was performed using TaqMan MicroRNA assays (Thermo Fisher Scientific 4427975 and 4440886).

### Immunoblotting

Cells were lysed in RIPA buffer supplemented with protease inhibitor cocktail (Roche 5056489001. 50 μg of protein were resolved by SDS-PAGE and transferred to PVDF membranes. Membranes were blocked with 5% milk in TBST, then incubated with primary antibodies overnight at 4°C, followed by 2 hours incubation with HRP-conjugated secondary antibodies (Thermo Fisher Scientific 31430; 31460) at rtp. Bands were visualized using SuperSignal™ West Pico PLUS Chemiluminescent Substrate (Thermo Fisher Scientific 34580) or Amersham™ ECL™ Select Western Blotting Detection Reagent (Cytiva RPN2235) and imaged on ChemiDoc MP.

For western blotting, the following antibodies were used: C15orf48 Antibody (Novus Biologicals NBP1-98391), NDUFA4 antibody (Abclonal A15693), Actin antibody (Santa Cruz sc-47778), GAPDH antibody (Proteintech 10494-1-AP).

### Flow cytometry

Cells were washed with PBS and stained with fluorophore-conjugated antibodies against surface antigens for 30 min at 4°C in the dark. After washing, cells designated for intracellular antigen detection were fixed and permeabilized using the eBioscience™ Foxp3 / Transcription Factor Staining Buffer Set (Thermo Fisher Scientific 00-5523-00) according to the manufacturer’s instructions, followed by staining with antibodies against intracellular antigens for 30 min at room temperature in the dark. Stained cells were washed in PBS and fixed in 2% PFA. The cells were analyzed on LSRFortessa^TM^ (BD Biosciences) and the data were analyzed using FlowJo (BD Biosciences).

For flow cytometry, the following antibodies were used: BD Horizon™ BUV395 Rat Anti-Mouse CD45 antibody (BD 564279), Brilliant Violet 711™ anti-mouse CD54 Antibody (Biolegend 116143), BD Pharmingen™ Alexa Fluor™ 488 Rat Anti-CD11b antibody (BD 557672), BD Pharmingen™ PerCP-Cy™5.5 Rat Anti-Mouse CD3 Molecular Complex antibody (BD 560527), BD Pharmingen™ PE-Cy™7 Rat Anti-Mouse CD25 antibody (BD 561780), BD Horizon™ BUV737 Rat Anti-Mouse CD19 (BD 612781), BD Horizon™ BV421 Hamster Anti-Mouse CD11c (BD 562782), BD Horizon™ BV605 Rat Anti-Mouse Ly-6C (BD 563011), BD Horizon™ BV650 Rat Anti-Mouse CD4 (BD 563232), BD Horizon™ BV786 Hamster Anti-Mouse CD69 (BD 564683), BD Pharmingen™ PerCP-Cy™5.5 Rat Anti-Mouse CD3 Molecular Complex (BD 560527), BD Horizon™ PE-CF594 Rat Anti-Mouse Ly-6G (BD 562700), BD Pharmingen™ PE-Cy™5 Rat Anti-Mouse CD8a (BD 561094), BD Horizon™ APC-R700 Rat Anti-Mouse F4/80 (BD 565787), BD Pharmingen™ APC-Cy™7 Rat Anti-Mouse Siglec-F (BD 565527). Dead cells were stained using Fixable Viability Stain 440UV (BD Biosciences 566332).

### XFe96 seahorse assay

Mitochondrial stress and glycolytic rate assays were performed using the Agilent Seahorse XFe/XFp Analyzer according to the manufacturer’s protocols. Monocytes, BMDMs and neutrophils were seeded at 100,000, 60,000 and 120,000 cells/well onto Seahorse assay plates, respectively. For oxidative burst assays, neutrophils were seeded onto plates pre-coated with 22.5 μg/mL Cell-Tak (Corning) to ensure adherence. The mitostress assay was conducted by sequential injection of 1 µM oligomycin, 1 µM FCCP, and 1 µM rotenone/antimycin A, while the glycolytic rate assay involved sequential injection of 1 µM rotenone/antimycin A and 50 mM 2-deoxyglucose injections. For oxidative burst measurements, respiration was inhibited in the cells first with 1 µM rotenone/antimycin A in the first injection, followed by stimulation by 100 nM PMA in the second injection. Measurements were taken for 2 hours, before 50 mM 2-deoxyglucose was injected. Extracellular acidification rate (ECAR) and oxygen consumption rate (OCR) were recorded every 3 minutes for all assays. These assays were done in mitostress media (XF basal DMEM (Agilent Technologies 10-357-5100) supplemented with 1 mM pyruvate (Sigma-Aldrich S8636), 2 mM glutamine (Life Technologies 25030081), and 10 mM glucose (Sigma-Aldrich G8644)). Data were analyzed using Agilent Seahorse Wave software.

To assess the activity of complex IV in cellular samples, the culture medium was replaced with MAS Buffer, composed of 70 mM sucrose, 220 mM mannitol, 10 mM potassium dihydrogen phosphate, 5 mM magnesium chloride, 2 mM HEPES, and 1 mM EGTA, adjusted to pH 7.2, and supplemented with 0.4% fatty acid-free BSA. A mixture containing N,N,N’,N’-tetramethyl-p-phenylenediamine (TMPD, Sigma-Aldrich T7394), ascorbate (Sigma-Aldrich A5960), ADP (Sigma-Aldrich A5285), FCCP (Sigma-Aldrich C2920), and digitonin was administered during the initial injection, resulting in final well concentrations of 0.5 mM TMPD, 2 mM ascorbate, 1 mM ADP, 1 μM FCCP, and 0.0025% digitonin. This is followed by 2 μM oligomycin, followed by 30mM potassium azide.

### Dissociation of lungs into single cells

Mice were euthanized by CO₂ asphyxiation, and the vessels in the lungs were perfused with PBS via the right ventricle prior to excision. Excised lung tissues were coarsely minced and resuspended in 2 mL of RPMI (HyClone SH30027.01) containing 0.4 mg/mL Liberase DL (Roche 5466202001), 20 U DNase I (Roche 4716728001), and 1% fatty acid-free BSA (Merck A8806). The tissue suspension was transferred to gentleMACSTM C Tubes (Miltenyi Biotec 130-093-237) and incubated at 37 °C for 30 min. The tissue was then mechanically dissociated on the gentleMACS Octo Dissociator (Miltenyi Biotec 130-096-427). Digestion was halted by the addition of ice-cold PBS containing 5% FBS and 1 mM EDTA. Cell suspensions were filtered through 70 µm strainers (SPL Life Sciences, 93070) and centrifuged at 500 × g for 10 min at 4°C. Erythrocytes were lysed using commercial lysis buffer (BioLegend 420301), and the resulting cell pellet was resuspended in FACS buffer (PBS with 2% FBS and 1 mM EDTA) for downstream flow cytometry or single-cell RNA sequencing.

### Single Cell RNA-seq

Single cell suspensions were obtained from lungs or bone marrow as described above. The cells from lungs were used directly, whilst the bone marrow cells were cultured with and without LPS for 4 hours. The cells were then stained with Biolegend TotalSeq™-B anti-mouse Hashtag antibodies. For the lung cells, wildtype and *C15orf48*-KO lung cells from untreated mice, and wildtype and *C15orf48*-KO lung cells from LPS-injected mice were stained with Hashtag 1 to 4, respectively (BioLegend 155831, 155833, 155835, 155837). For bone marrow cells, untreated wildtype and *C15orf48*-KO cells, and LPS-treated wildtype and *C15orf48*-KO cells were stained with Hashtag 5-8, respectively (BioLegend 155839, 155841, 155843, 155845). Briefly, 1 million cells were stained in 100 μL of 0.2% FCS/PBS containing 0.5 μL of the hashtag for 30 min on ice. The cells were then washed 1 time with 3 mL of 0.2% FCS/PBS, pelleted at 300 rcf for 5 min at 4 °C and resuspended in 100 μL of 0.2% FCS/PBS containing 1:1000 anti-rat-Alexa Fluor™ 647 (Thermo Fisher Scientific A48272). The cells were incubated on ice for another 15 min, before they were washed and resuspended in 500 μL of 0.2% FCS/PBS containing 5 μL of 7-AAD Viability Staining Solution (BioLegend 420404). 200,000 cells from each sample were sorted by flow cytometry to select for live and stained cells. The cells were then pelleted and resuspended in 0.2% FCS/PBS to be counted. Equal number of cells from each sample were combined and strained through 40 μm Flowmi® Cell Strainers (Bel-Art™, BAH136800040). Library preparation were prepare using Chromium Next GEM Single Cell 3’ Reagent kit v3.1 (10x Genomics, 1000269) workflow according to the manufacturer’s instructions.

### Single-cell RNA-seq preprocessing and demultiplexing

Raw sequencing data (FASTQ files) were processed using Cell Ranger version 6 (10x Genomics) with default settings to produce gene-by-cell count matrices. Hashtag oligonucleotide (HTO) data from multiplexed samples were used for sample identification, assigning each cell to its source condition through the HTODemux function in Seurat. Cells identified as doublets or those with ambiguous HTO assignments were excluded from further analysis. The filtered count matrices were then analysed using Seurat v4, following the package’s standard workflow. For quality control, cells were retained if they had between 200 and 6,000 detected genes (nFeature_RNA) and fewer than 15% of transcripts mapping to mitochondrial genes, to remove doublets, empty droplets, and low-quality cells. After QC, gene expression data were normalized and log-transformed in Seurat prior to downstream analyses and visualization. Cell types were annotated using Seurat’s label transfer approach (TransferData function).

### Differential expression and GSEA pathway enrichment analysis

For each identified cell type, differential gene expression (DGE) analysis between KO-LPS and WT-LPS groups was conducted using the FindMarkers function in Seurat. Only genes detected in at least 5% of cells were included, with no minimum log fold change cutoff applied (min.pct = 0.05, logfc.threshold = 0). The ranked list based on log₂ fold changes was then used as input for gene set enrichment analysis (GSEA), performed with the fgsea package. Gene sets for Gene Ontology Biological Process (GO:BP) terms were sourced from the C5 collection of MSigDB using the msigdbr package. Pathways selected for analysis contained between 5 and 500 genes. Normalized enrichment scores (NES) were calculated, and only pathways with an adjusted p-value below 0.5 were visualized. The most significant pathways by NES magnitude were selected, and hierarchical clustering was applied to the rows (genes) for heatmap generation with the pheatmap package.

## Data availability

The scRNA-seq dataset on lungs from LPS-injected WT and *C15orf48*-KO mice is available under GSE339377. The scRNA-seq dataset on Kp-infected cells was taken from NCBI Gene Expression Omnibus repository with the accession number GSE184290^8^. scRNA-seq data on PBMCs treated with LPS and Pam3CSK4 was taken from Broad Institute Single Cell Portal (SCP550)^9^.

## Statistical Analysis

Two-tailed unpaired Student’s t test, one- or two-way ANOVA and Gehan-Breslow-Wilcoxon test were calculated using GraphPad Prism. ANCOVA and Benjamini-Hochberg procedure were calculated using R^7^. All the graphs were presented as mean ± standard error.

