## Supplemental data for "Complex IV Subunit C15orf48 Integrates Electron Transport Chain Function with Amino Acid and Lipid Metabolism to Regulate Innate Immunity"

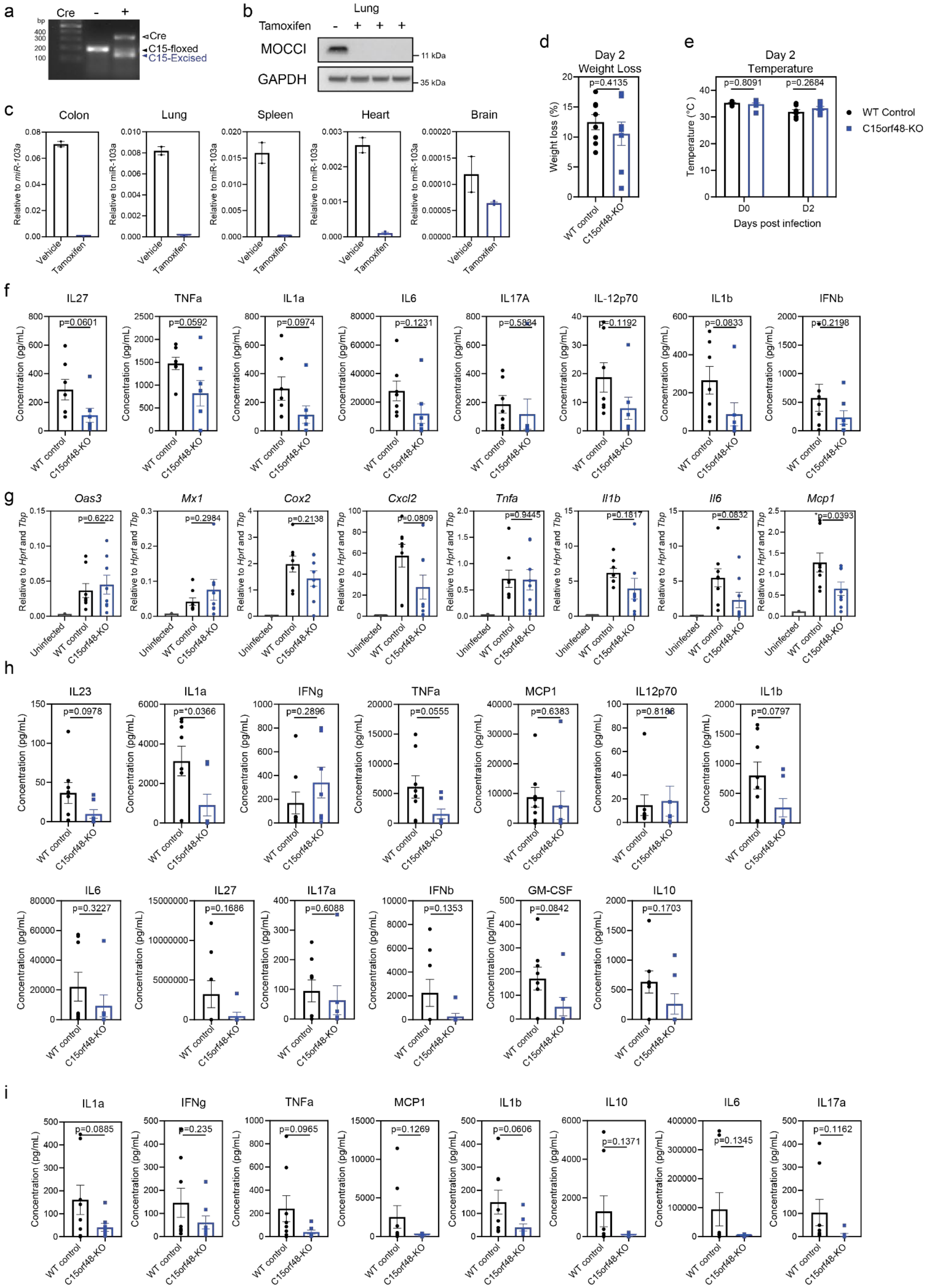

a
Cre
-
+
bp
400
300
200
100
Cre
C15-floxed
C15-Excised
b
Lung
Tamoxifen
MOCCI
GAPDH
11 kDa
35 kDa
c
Colon
Lung
Spleen
Heart
Brain
Relative to miR-103a
Vehicle
Tamoxifen
d
Day 2
Weight Loss
p=0.4135
Weight loss (%)
WT control
C15orf48-KO
e
Day 2
Temperature
p=0.8091
p=0.2684
Temperature (°C )
D0
D2
Days post infection
WT Control
C15orf48-KO
f
IL27
TNFa
IL1a
IL6
IL17A
IL-12p70
IL1b
IFNb
p=0.0601
p=0.0592
p=0.0974
p=0.1231
p=0.5834
p=0.1192
p=0.0833
p=0.2198
Concentration (pg/mL)
WT control
C15orf48-KO
g
Oas3
Mx1
Cox2
Cxcl2
Tnfa
Il1b
Il6
Mcp1
p=0.6222
p=0.2984
p=0.2138
p=0.0809
p=0.9445
p=0.1817
p=0.0832
p=0.0393
Relative to Hprt and Tbp
Uninfected
WT control
C15orf48-KO
h
IL23
IL1a
IFNg
TNFa
MCP1
IL12p70
IL1b
p=0.0978
p=0.0366
p=0.2896
p=0.0555
p=0.6383
p=0.8186
p=0.0797
IL6
IL27
IL17a
IFNb
GM-CSF
IL10
p=0.3227
p=0.1686
p=0.6088
p=0.1353
p=0.0842
p=0.1703
Concentration (pg/mL)
WT control
C15orf48-KO
i
IL1a
IFNg
TNFa
MCP1
IL1b
IL10
IL6
IL17a
p=0.0885
p=0.235
p=0.0965
p=0.1269
p=0.0606
p=0.1371
p=0.1345
p=0.1162
Concentration (pg/mL)
WT control
C15orf48-KO

**Supplementary Figure 1. *C15orf48*-KO mice are less susceptible to Kp infection.**

(a) Genotyping gel for ROSA-creERT2 (cre) positive and negative *C15orf48*-flox/flox mice after tamoxifen administration.

(b) MOCCI expression in lung after tamoxifen administration and LPS injection. (n = 3)

(c) MiR-147b expression in various organs with and without tamoxifen administration. (n=2 without tamoxifen, 3 with tamoxifen administration)

(d) Percentage weight loss of WT and *C15orf48*-KO mice two days after Kp infection. P = two-tailed unpaired Student's t test; n = 8 biological replicates over two repeats.

(e) Body temperature of WT and *C15orf48*-KO mice before and two days after Kp infection. P = two-way ANOVA; n = 8 biological replicates over two repeats.

(f) Cytokine analysis of BALF taken from WT or *C15orf48*-KO mice two days after Kp infection. P = two-tailed unpaired Student's t test; n = 8 biological replicates over two repeats.

(g) qPCR of lungs from WT or *C15orf48*-KO mice two days after Kp infection. P = two-tailed unpaired Student's t test between WT and *C15orf48*-KO mice; n = 8 biological replicates over two repeats.

(h) Cytokine analysis of BALF taken from WT or *C15orf48*-KO mice two days after Kp infection. P = two-tailed unpaired Student's t test; n = 8 biological replicates over two repeats.

(i) Cytokine analysis of plasma taken from WT or *C15orf48*-KO mice two days after Kp infection. P = two-tailed unpaired Student's t test; n = 8 biological replicates over two repeats.

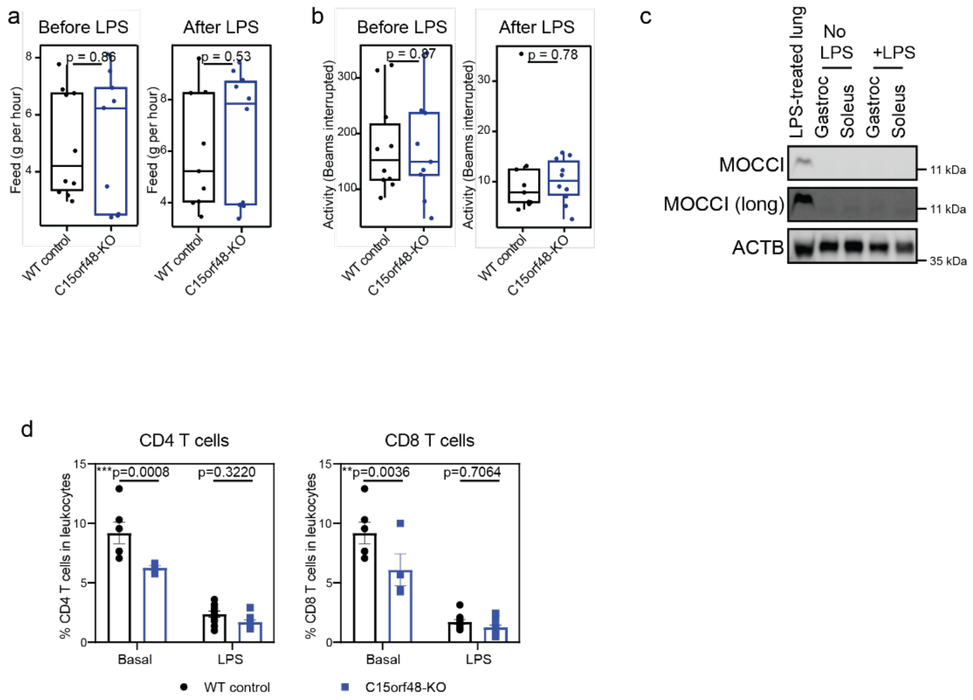


**Supplementary Figure 2.** ***C15orf48*-KO mice exhibit lower systemic inflammation during endotoxemia.**

(a) Quantification of the amount of food intake before and after LPS injection of WT and *C15orf48*-KO mice by the PhenoMaster. P = two-way ANCOVA; n = 10 biological replicates over two repeats.

(b) Quantification of the activity before and after LPS injection of WT and *C15orf48*-KO mice by the PhenoMaster. P = two-way ANCOVA; n = 10 biological replicates over two repeats.

(c) Expression of MOCCI in skeletal muscle before and after LPS injection. LPS-treated lung was included as positive control. n = 3.

(d) Flow cytometry analysis of infiltrated T cells in lungs from mice injected with PBS or LPS. P = two-way ANOVA; n = 6 and 12 biological replicates for basal and after LPS, respectively, over three repeats.

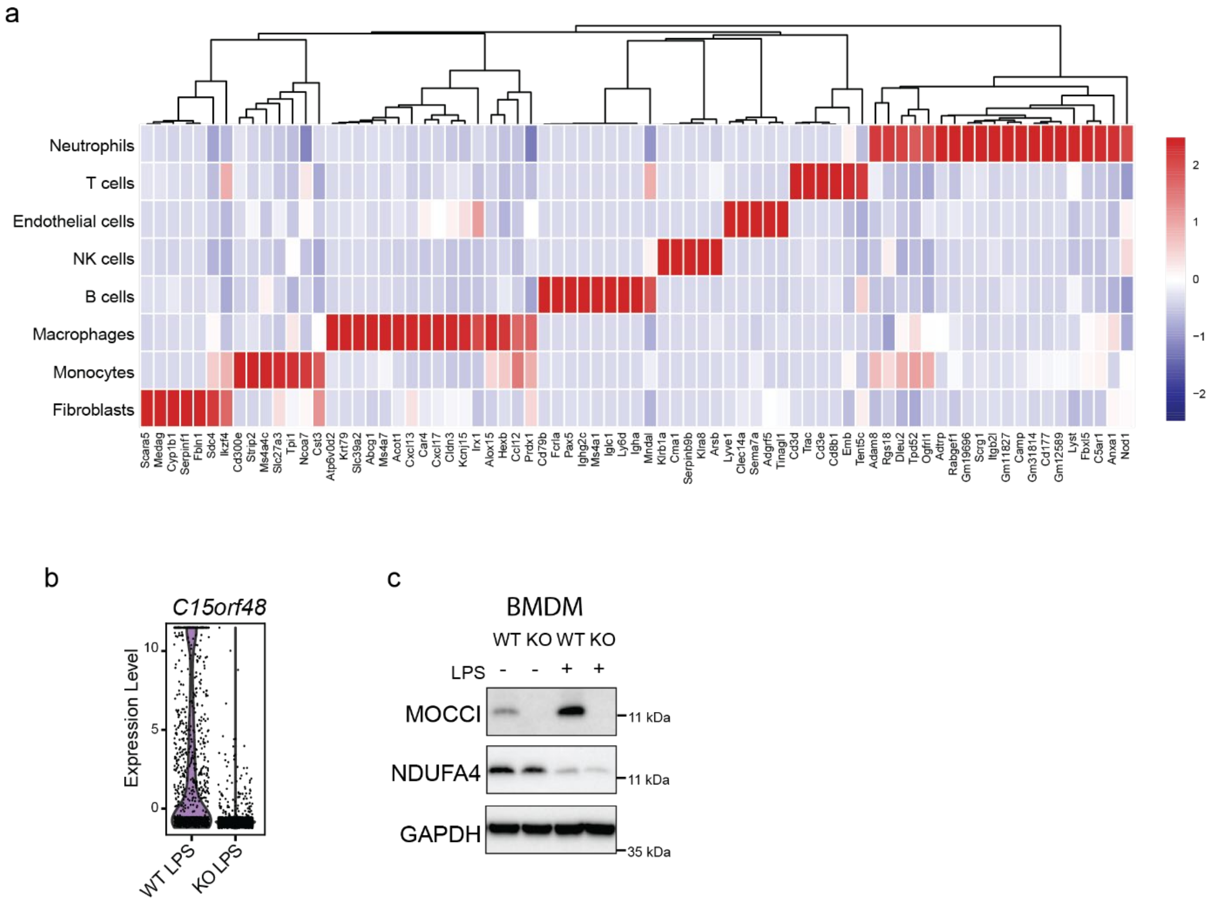


**Supplementary Figure 3. *C15orf48* is expressed in human and mouse monocytes.**

(a) Identification of cell types in the scRNA-seq done on single cell isolates from lungs, based on the expression of marker genes. X-axis contains different genes while Y-axis clusters cells with similar gene expression together. Cell identity is predicted based on their expression of marker genes.

(b) Violin plot of *C15orf48* expression in WT and *C15orf48*-KO mice after LPS injection.

(c) Western blot of MOCCI and NDUFA4 in WT and *C15orf48*-KO murine bone marrow-derived macrophages after 16 hours treatment with PBS and LPS. n = 3 biological repeats.

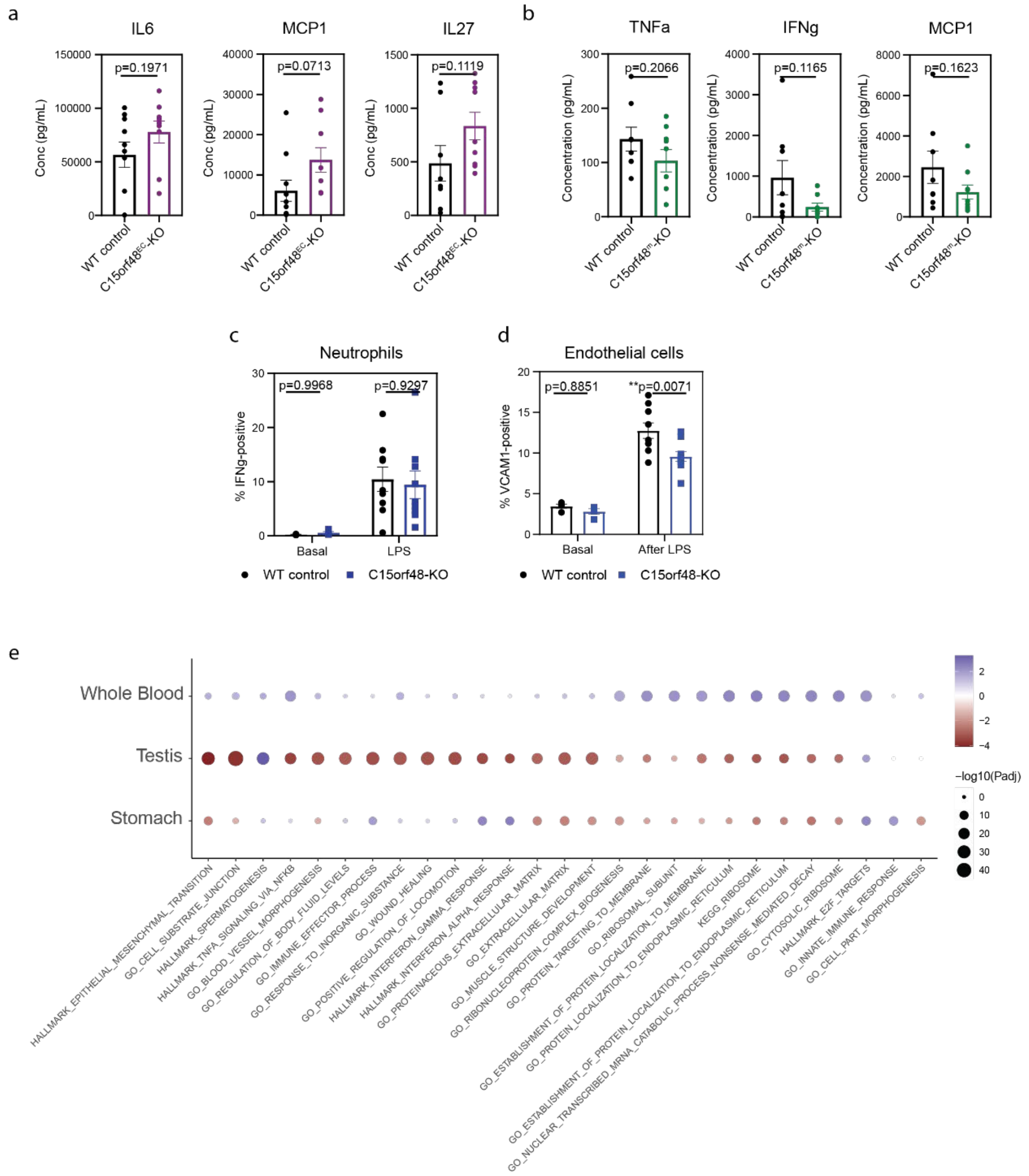


**Supplementary Figure 4. *C15orf48* plays opposing roles in myeloid and endothelial cells during LPS-induced systemic inflammation.**

(a) Levels of proinflammatory cytokines in plasma of WT and *C15orf48*$^{EC}$-KO mice one day after LPS injection. P = two-tailed unpaired Student's t test; n = 9 biological replicates over two repeats.

(b) Levels of proinflammatory cytokines in plasma of WT and *C15orf48*$^{m}$-KO mice one day after LPS injection. P = two-tailed unpaired Student's t test; n = 8 biological replicates over two repeats.

(c) Percentage of IFNγ- positive neutrophils in lungs from WT and *C15orf48*-KO mice at basal and after LPS injection. P = two-way ANOVA; n = 6 biological replicates for mice at basal and 9 for LPS-injected mice from three repeats.

(d) Percentage of VCAM1-positive endothelial cells in lungs from WT and *C15orf48*-KO mice at basal and after LPS injection. P = two-way ANOVA; n = 6 biological replicates for mice at basal and 9 for LPS-injected mice from three repeats.

(e) Correlation of *C15orf48* expression with each Hallmark in whole blood, testis and stomach.

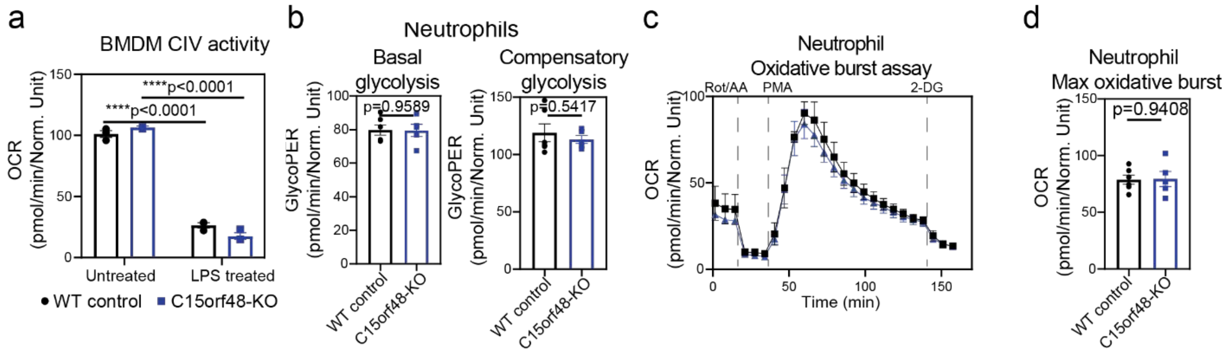


**Supplementary Figure 5. *C15orf48* deficiency alters bioenergetics, redox balance, and cytokine secretion in monocytes.**

(a) Complex IV activity of murine BMDMs at basal and after 6h LPS treatment. P = two-way ANOVA; n = 3 biological replicates.

(b) Basal and compensatory glycolytic rate of WT and *C15orf48*-KO neutrophils at basal, as measured by Seahorse XF Glycolytic Rate assay. P = two-tailed unpaired Student's t test; n = 6 biological replicates over two repeats.

(c) Oxidative activity in WT and *C15orf48*-KO neutrophils was measured by Seahorse XF in an oxidative burst assay. P = two-tailed unpaired Student's t test; n = 6 biological replicates over two repeats.

(d) Quantification of the maximum oxidative burst in WT and *C15orf48*-KO neutrophils, was measured by the oxidative burst assay in (e). P = two-tailed unpaired Student's t test; n = 6 biological replicates over two repeats.

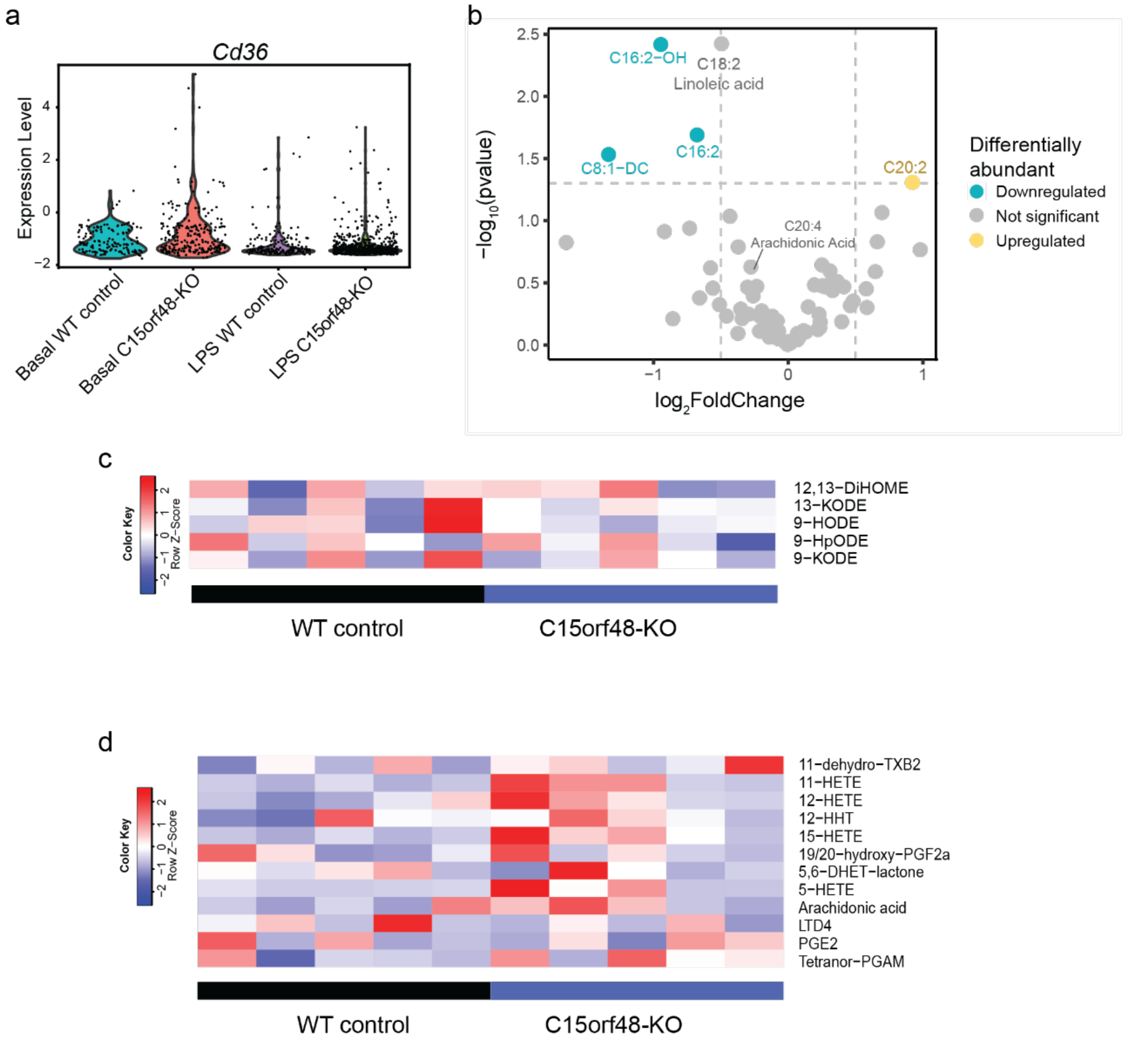


**Supplementary Figure 6. Reduced FAO in *C15orf48*-KO monocytes is associated with reduced monocyte activation.**

(a) Expression of *Cd36* in monocytes in lungs, based on scRNA-seq done on cells isolated from lungs.

(b) Targeted metabolomics probing for acylcarnitines on plasma from WT and *C15orf48*-KO mice. $Log_2$ fold change (*C15orf48*-KO/WT) for each detected metabolite are plotted. n = 4 biological replicates. P-adjusted is calculated by the Benjamini-Hochberg procedure.

(c) Relative levels of linoleic acid derivatives measured from targeted lipid mediator metabolomics on WT and *C15orf48*-KO monocytes at basal. n = 5 biological replicates. Each biological replicate contained cells pooled from 3-4 mice.

(d) Relative levels of arachidonic acid derivatives measured from targeted lipid mediator metabolomics on WT and *C15orf48*-KO monocytes at basal. n = 5 biological replicates. Each biological replicate contained cells pooled from 3-4 mice.

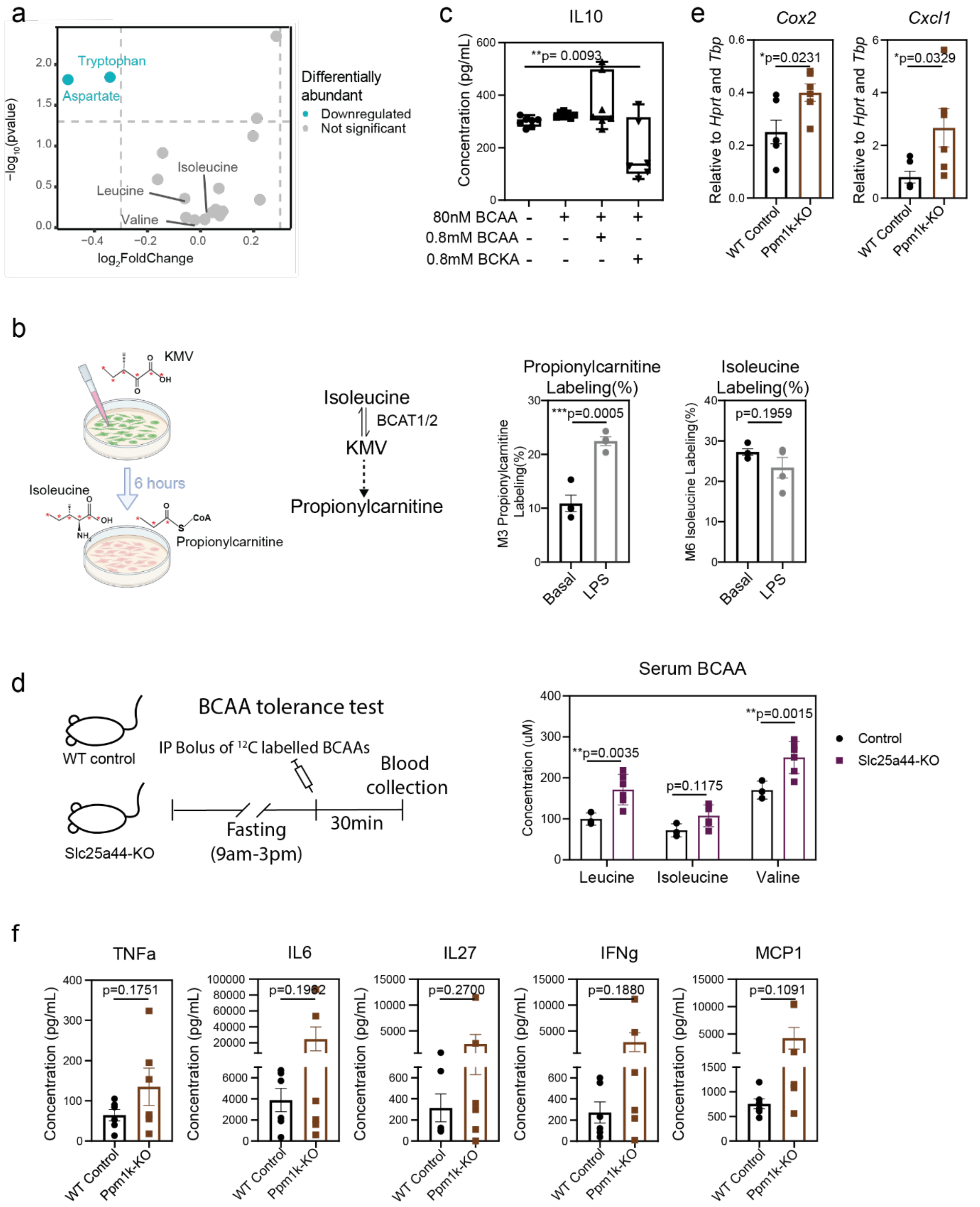


**Supplementary Figure 7. Reduced BCAA metabolism in *C15orf48*-KO monocytes is associated with reduced monocyte activation.**

(a) Targeted metabolomics probing for amino acids on plasma from WT and *C15orf48*-KO mice. $Log_2$ fold change (*C15orf48*-KO/WT) for each detected metabolite are plotted. n = 4 biological replicates. P-adjusted is calculated by the Benjamini-Hochberg procedure.

(b) Measured propionylcarnitine and isoleucine isotopologue labelling in WT and *C15orf48*-KO monocytes, treated with 1 mM [$^{13}C_6$] alpha-keto-beta-methyl-n-valeric acid (KMV).

(c) Cytokine secretion after LPS from WT monocytes cultured in 80 nM BCAA with 80 mM BCAA or 80 mM BCKA. P = one-way ANOVA; n = 6 biological replicates per condition from two repeats.

(d) (Left) Experimental design of the BCAA tolerance test. WT control and *Slc25a44*-KO mice were fasted from 9:00 AM to 3:00 PM, followed by intraperitoneal injection of a BCAA mixture containing $^{12}$C-leucine, $^{12}$C-isoleucine, and $^{12}$C-valine. Blood samples were collected 30 min after injection for metabolomic analysis. (Right) Serum concentrations of BCAAs after the BCAA tolerance test. P = two-way ANOVA. n = 3 WT and 6 *Slc25a44*-KO biological replicates.

(e) qPCR of proinflammatory genes in lungs from WT and *Ppm1k*-KO mice after LPS injection. P = two-tailed unpaired Student's t test; n = 6 biological replicates over two repeats.

(f) Quantification of cytokines in plasma from WT and *Ppm1k* -KO mice after LPS injection. P = two-tailed unpaired Student's t test; n = 6 biological replicates over two repeats.
